# Characterising cancer cell responses to cyclic hypoxia using mathematical modelling

**DOI:** 10.1101/2024.06.25.600569

**Authors:** Giulia Celora, Ruby Nixson, Joe Pitt-Francis, Philip Maini, Helen Byrne

**Author notes:** These authors contributed equally to this work (Joe Pitt-Francis, Philip Maini, Helen Byrne).

## Abstract

In vivo observations show that oxygen levels in tumours can fluctuate on fast and slow timescales. As a result, cancer cells can be periodically exposed to pathologically low oxygen levels; a phenomenon known as cyclic hypoxia. Yet, little is known about the response and adaptation of cancer cells to cyclic, rather than, constant hypoxia. Further, existing in vitro models of cyclic hypoxia fail to capture the complex and heterogeneous oxygen dynamics of tumours growing *in vivo*. Mathematical models can help to overcome current experimental limitations and, in so doing, offer new insights into the biology of tumour cyclic hypoxia by predicting cell responses to a wide range of cyclic dynamics. We develop an individual-based model to investigate how cell cycle progression and cell fate determination of cancer cells are altered following exposure to cyclic hypoxia. Our model can simulate standard *in vitro* experiments, such as clonogenic assays and cell cycle experiments, allowing for efficient screening of cell responses under a wide range of cyclic hypoxia conditions. Simulation results show that the same cell line can exhibit markedly different responses to cyclic hypoxia depending on the dynamics of the oxygen fluctuations. We also use our model to investigate the impact of changes to cell cycle checkpoint activation and damage repair on cell responses to cyclic hypoxia. Our simulations suggest that cyclic hypoxia can promote heterogeneity in cellular damage repair activity within vascular tumours.

## 1 Introduction

Uncontrolled proliferation is one of the hallmarks of cancer (Hanahan, 2022). However, experimental evidence shows that intra-tumour heterogeneity in proliferation activity is a leading cause of treatment failure, with small numbers of quiescent (*i.e*., non-proliferative) cancer cells driving drug resistance and relapse (Aguirre-Ghiso, 2007; Tomasin and Bruni-Cardoso, 2022). This observation highlights the need to understand what environmental and subcellular signals regulate quiescence in cancer (Tomasin and Bruni-Cardoso, 2022).

The mitotic cell cycle is commonly divided into four phases: G1 (growth in preparation for DNA replication), S (DNA synthesis), G2 (growth and preparation for mitosis) and M (mitosis). As cells proceed through the cell cycle there are two key decisions to make: whether to initiate DNA replication and whether to undergo mitosis. These decisions are regulated by integrating multiple cellular signalling pathways and extracellular stimuli. At the cell scale, control mechanisms (or checkpoints) guarantee timely and accurate replication of the genome (in the S phase) and its correct segregation into two daughter cells (in the M phase). At the tissue scale, environmental cues, such as growth factors, nutrient levels and mechanical stress, can favour re-entry into, or arrest of, the mitotic cycle to maintain tissue homeostasis by regulating check-point dynamics. To maintain high rates of proliferation, cancer cells must disrupt cell cycle regulation mechanisms designed to prevent the replication of damaged/neoplastic cells. Such behaviour is usually associated with mutations or dysregulation of proteins that control cell cycle checkpoints; specifically, cell cycle control in response to DNA damage and S-phase entry (Matthews et al, 2022). Nonetheless, cells may still benefit from having functioning checkpoints that induce quiescence and enable cancer cell survival under unfavourable conditions.

As a solid tumour develops, excessive cell proliferation leads to an imbalance between oxygen supply and demand, resulting in pathologically low oxygen levels (*i.e*., hypoxia) at distance from the vasculature. Hypoxia is a known driver of cellular quiescence and has been associated with poor treatment outcomes. As hypoxia is toxic for proliferating cells, particularly those actively synthesising DNA, cells that reside in hypoxic regions may enter a quiescent state. By transiently exiting the cell cycle, these cells are able to withstand adverse environmental conditions.

Oxygen levels in vascularised tumours are both spatially and temporally heterogeneous (Kawai et al, 2022; Matsumoto et al, 2010). As a result, regions in which cells are periodically exposed to hypoxia can arise, a phenomenon known as *cyclic hypoxia*. While constant hypoxia has been widely studied, relatively little is known about how cancer cells respond to fluctuating oxygen levels and how cyclic hypoxia contributes to cancer cell proliferation. While constant hypoxia is known to induce cell quiescence (Höckel, M and Vaupel, P, 2001), the interplay between cell cycle progression and cell survival under fluctuating oxygen environments remains to be elucidated.

There are several experimental challenges associated with quantifying cyclic hypoxia *in vivo* and replicating such conditions *in vitro*. Indeed, a key limitation in studying cyclic hypoxia is the development of experimental models that replicate the oxygen dynamics experienced by tumours growing *in vivo*. While constant hypoxia typically affects tumour regions at a significant distance from vessels, cyclic hypoxia is observed both close to, and far from, blood vessels, with periods ranging from seconds to days (Bader et al, 2021a; Ron et al, 2019). Furthermore, there can be large variability in the amplitude and frequency of oxygen fluctuations within the same tumour. High-frequency fluctuations are usually associated with vasomotor activity, while vascular remodelling and treatment, can generate cycles with longer periods (Michiels et al, 2016). Recent work suggests that self-sustained fluctuations in blood flow might be related to the topology of blood vessel networks, which is known to be abnormal in tumours (Ben-Ami et al, 2022). In previous work (Celora et al, 2022; Celora, 2022), we have shown how mathematical modelling can be combined with experimental data to study the impact of short-term exposure to a wide range of cyclic hypoxia protocols on cell cycle progression in the colorectal RKO cancer cell line. Here, we extend our cell cycle model to investigate the long-term impact of time-varying oxygen levels on cancer cell survival and the emergent population growth dynamics. The flexibility of our modelling framework allows us to investigate how cell cycle checkpoint and damage repair signalling influence cancer cells’ adaptation to different forms of cyclic hypoxia. In doing so, we obtain new insight into how cyclic hypoxia may contribute to intra-tumour heterogeneity and treatment resistance by favouring the selection of cancer cells which differ in their ability to repair damage.

The paper is organised as follows. In Section 2, we review what is currently known about cell cycle progression and cell survival in different hypoxic environments. In Section 3, we present a stochastic, individual-based (IB) model of the cell cycle in hypoxia which aims to capture aspects of the biology presented in Section 2. In Section 4.1, we validate our model by simulating growth dynamics in constant environmental conditions and comparing model output with experimental observations. In Section 4.2, we use our model to study how different fluctuating hypoxic environments affect the growth dynamics (see Section 4.2.1) and survival outcomes (see Section 4.2.2) of cancer cell populations. In Section 4.3, simulations of serial passage experiments reveal how alterations to damage repair and cell cycle checkpoint signalling may affect cancer cell responses and adaptation to cyclic hypoxia. In Section 5, we explain how our results increase our understanding of how cyclic hypoxia may contribute to tumour heterogeneity by allowing the coexistence of cells with different levels of damage repair capacity. We conclude in Section 6 by summarising our results and outlining possible directions for future research.

## 2 Cell (dys-)regulation in hypoxia

When characterising cell responses to hypoxia, it is important to account for the oxygen concentration to which the cells are exposed. In this study, we use the term “hypoxia” to refer to oxygen levels below *c*_*H*_ = 1% O_2_, which is often referred to as pathological hypoxia (McKeown, 2014). In practice, a tumour’s tolerance to oxygen shortages depends on its tissue of origin. As such, this threshold should be viewed as an upper bound, rather than an absolute value (McKeown, 2014).

The schematic in Figure 1 summarises how prolonged exposure to hypoxia affects cell physiology by disrupting two fundamental processes: DNA synthesis and repair. The consequences of these perturbations are cell cycle phase specific (see Figure 1b).

**Fig. 1:**
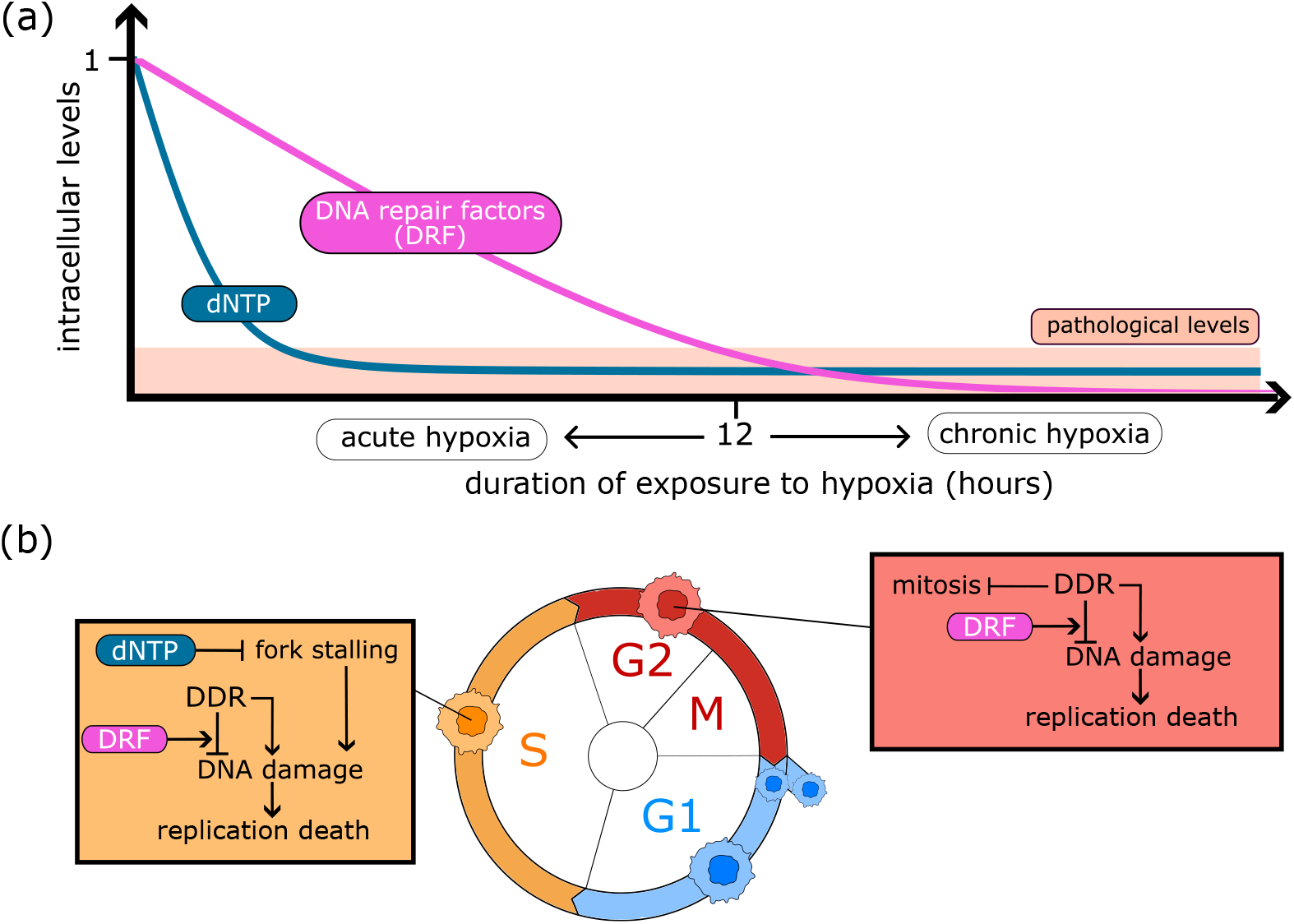
Schematic representation of cell responses as a function of hypoxia duration. (a) Intracellular levels of dNTP quickly drop to pathological levels in hypoxia, determining cell responses to acute hypoxia. Levels of DNA repair factors decrease more slowly than dNTP and, hence, drive cell responses to chronic hypoxia. (b) Cell cycle-specific role of dNTP and DRF levels on the regulation of intracellular mechanisms: DNA synthesis and damage accumulation/repair. Keys: arrow-heads indicate stimulation; bar-heads indicate inhibition; DDR denotes *DNA damage response* and dNTP denotes *deoxynucleotide triphosphates*. More details are given in the main text.

*In vitro* experiments have shown the rapid reduction in the initiation and progression of DNA synthesis in cells exposed to hypoxia (Foskolou et al, 2017; Pires et al, 2010b). This behaviour has been attributed to impaired functioning of the enzyme ribonucleotide reductase (RNR) (Foskolou et al, 2017; Olcina et al, 2010), which mediates *de novo* production of *deoxynucleotide triphosphates* (dNTPs). Since dNTPs are the building blocks of DNA, the reduction in dNTP levels prevents cells from initiating DNA synthesis (arrest in the G1 phase) and causes DNA synthesis to stall (arrest in the S phase). The stalling of DNA synthesis activates the DNA damage response (DDR), stabilising open replication forks and allowing cells in the S-phase to withstand replication stress. However, exposure to hypoxia also activates an energy-preserving program (Pires et al, 2010a) resulting in reduced production of DNA repair factors (DRF) and, hence, reduced ability to stabilise stalled replication forks. If hypoxic conditions are prolonged, more than 12 hours, arrest in the S phase becomes irreversible and leads eventually to cell death (Pires et al, 2010b; Ng et al, 2018). By contrast, cells that arrest before initiating DNA synthesis can tolerate prolonged exposure to hypoxia since they are not sensitive to replication stress. Differences in the time scales associated with the decreases in levels of dNTPs and DNA repair factors enable cells to distinguish between acute (less than 12 hours) and chronic (more than 12 hours) hypoxia. As a result, cells can adapt their response to oxygen dynamics rather than responding instantaneously to changes in oxygen levels.

If oxygen levels are restored after acute exposure to hypoxia, cells in the S phase can resume DNA synthesis although they may accumulate additional damage during re-oxygenation (Bader et al, 2021b). Depending on the amount of stress/damage sustained, activation of DDR signalling may cause these cells to accumulate in the G2 phase and prevent them from entering mitosis (Bristow and Hill, 2008; Goto et al, 2015; Olcina et al, 2010). Damaged cells that successfully repair any damage they have accumulated eventually enter mitosis and replicate; otherwise, they undergo reproductive death (either via activation of the senescence program or via cell death). Regulation of the DDR signalling and damage repair is therefore crucial in determining the longterm impact of hypoxia on cancer cell responses; conversely, hypoxia is known to shift the damage repair capacity of cells (Begg and Tavassoli, 2020).

In our previous work (Celora et al, 2022), we focussed on modelling cell responses to acute exposure to constant and cyclic hypoxia. As such, we neglected the role of DNA repair factors and the impact of hypoxia on cell viability. Here, we show how these effects can be included in our framework.

## 3 An individual-based model of *in vitro* cancer cell dynamics in hypoxia

### 3.1 Model overview

We consider a population of cells that are in a well-mixed (*i.e*., spatially homogeneous) environment and exposed to externally prescribed, time-varying oxygen levels, *c* = *c*(*t*) [O_2_%]. This mimics typical cell culture experiments in oxygen chambers (Kim et al, 2021). For simplicity, we focus on the early stages of population growth, when competition for space and nutrients can be neglected.

We represent each cell as an individual which can profilerate or die with probabilities that depend on their state. Each cell is characterised by five state variables (see Table 1). The categorical variable *z* indicates the position along the cell cycle (cell cycle state), while the four continuous state variables describe respectively DNA content (*x*), damage levels (*y*), intracellular levels of dNTP (*m*_dNTP_) and DNA repair factors (*m*_DRF_). Continuous state variables are included to account for the dynamics of intracellular processes that regulate cell proliferation (via progression through the cell cycle) and death in oxygen-fluctuating environments. Variables *x* and *m*_dNTP_ are introduced to describe the evolution of DNA synthesis in the S phase; variables *y* and *m*_DRF_ are introduced to describe the processes of damage repair.

**Table 1:** List of the variables characterising a cell (individual) state with a brief description and the range of values that they can take. The notation (a.u.) stands for arbitrary units.

| Variable | Description | Values |
| --- | --- | --- |
| $m_{\text{dNTP}}^{(i)}(t)$ | intracellular dNTP levels in cell $i$ at time $t$ (a.u.) | $[0, 1]$ |
| $m_{\text{DRF}}^{(i)}(t)$ | intracellular DRF levels in cell $i$ at time $t$ (a.u.) | $[0, 1]$ |
| $x^{(i)}(t)$ | DNA content in cell $i$ at time $t$ | $[1, 2]$ |
| $y^{(i)}(t)$ | damage/stress level in cell $i$ at time $t$ (a.u.) | $[0, \infty)$ |
| $z^{(i)}(t)$ | cell cycle state of cell $i$ at time $t$ | $\{G_1, C_1, S, G_2, C_2\}$ |

Proliferation, death and state changes of each cancer cell are described by time-discrete stochastic processes. We consider discrete time points: *t*_*n*_ = *n*Δ*t* ∈ [0, *t*_*f*_], where *t*_*f*_ is the final time of the simulations and the time-step Δ*t* ∈ R^+^ is chosen to be sufficiently small to resolve all dynamic processes included in the model. The flow chart in Figure 2 summarises the procedure used at each time-step to simulate cell fate decisions (*i.e*., death, division or progression along the cell cycle) and intracellular processes (*i.e*., DNA replication and damage repair). In the rest of this Section, we first briefly describe the rules used to simulate cell fate decisions; we then outline the rules used to simulate intracellular processes; namely, DNA synthesis, DNA repair and dNTP and DRF production. To conclude, we summarise the IB simulations we perform and how they are initialised.

**Fig. 2:**
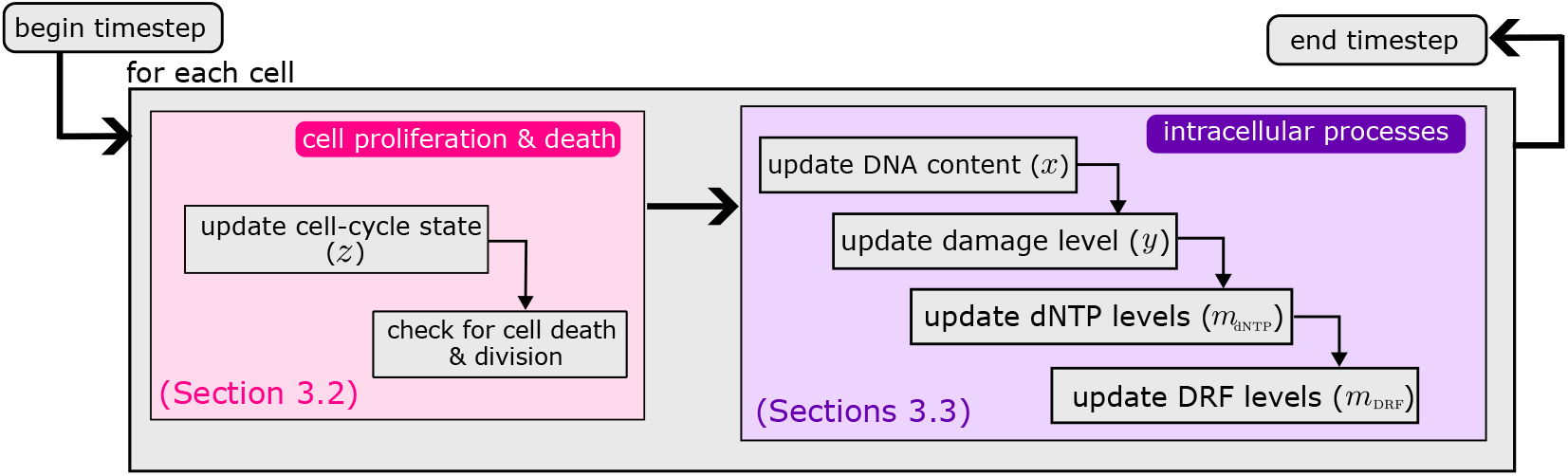
Flowchart illustrating how we implement our stochastic IB model to simulate *in vitro* cancer cell dynamics in hypoxia. The algorithm comprises two main subroutines: simulation of cell proliferation and death (pink shaded area; and simulation of intracellular processes (purple shaded area; Details given in Appendix A)).

### 3.2 Modelling cell proliferation and death

Following (Celora et al, 2022), we assume that cells exist in one of five cell cycle states: *G*_1_, *C*_1_, *S, G*_2_, *C*_2_. Table 2 summarises the role of each cell cycle state in the model and how they map to (biological) cell cycle phases. At any time-step *t*_*n*_, cells can update their cell cycle state, divide or die with probabilities that depend on the values of their state variables (see Table 1) and oxygen levels as summarised in the schematic in Figure 3.

**Table 2:** Description of the cell cycle states included in our model and how they map to the standard biological cell cycle phases: G1, S, G2 and M.

| Cell cycle state ($z$) | Description | Cell cycle phase |
| --- | --- | --- |
| $G_1$ | cells preparing to initiate DNA replication | G1 |
| $C_1$ | cells ready to start DNA replication but arrested due to checkpoint activation | G1 |
| $S$ | cells replicating DNA | S |
| $G_2$ | cells that have completed DNA replication and are preparing for cell division | G2/M |
| $C_2$ | cells that have completed DNA replication but can not enter mitosis because of checkpoint activation | G2 |

**Fig. 3:**
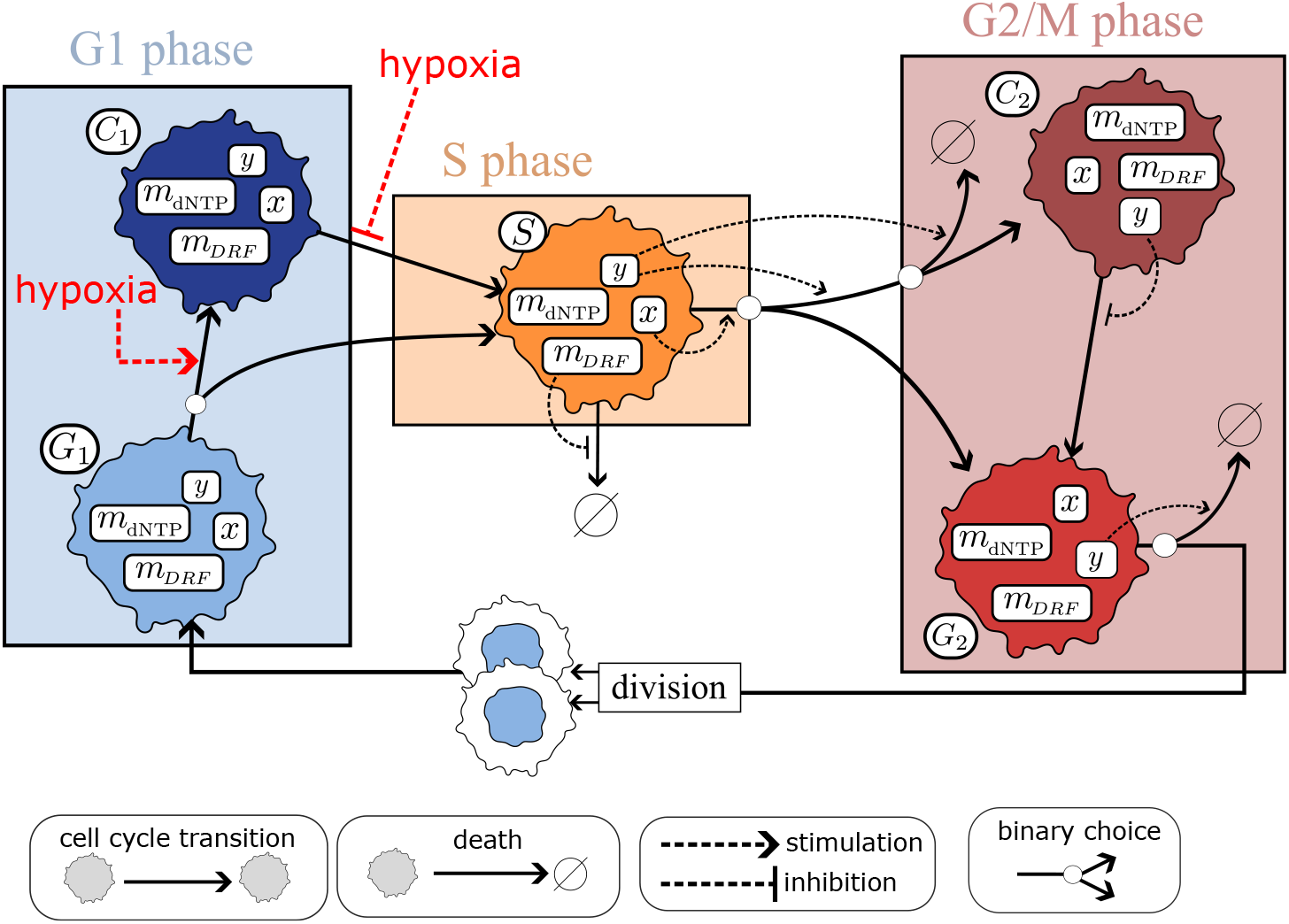
Description of our cell cycle model. Cells can exist in one of 5 cell cycle states (*z* ∈ {*G*_1_, *C*_1_, *S, G*_2_, *C*_2_}). Transitions between cell cycle states and cell death depend upon a cell state or oxygen levels as detailed in Section A. We illustrate how the internal variables regulate progression through the cell cycle. Note that we indicate only the internal variables that regulate cell cycle transitions. At the points where the continuous arrows bifurcate, only one of the possible paths is chosen. The symbol ∅ indicates loss of replication capacity, either via cell death and/or senescence.

We model the stimulatory/inhibitory effects of intracellular and environmental (*i.e*., oxygen) factors on cell cycle transitions by using the sigmoid function

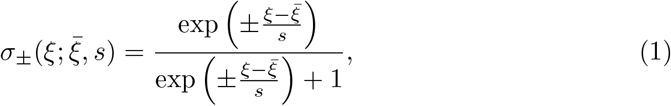

which is commonly used in modelling non-linear activation responses that are mediated by multistep processes (Ferrell et al, 2011). In Eq. (1) the subscript indicates whether the variable *ξ* induces a stimulatory (+) or inhibitory (−) effect. As shown in Figure 4, the parameter 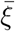 shifts the sigmoid function so that its inflection point is located at 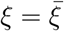, while the parameter *s* regulates the steepness of the sigmoidal curve. For *s* → 0, *σ*_*±*_ converges to a Heaviside step function (switch-like response), while larger values of *s* correspond to a smoother, graded response. Given this formalism, we translate the diagram in Figure 3 into a set of rules that determine cell death, cell division and how the cell cycle state of each cell is updated from time *t*_*n*_ to time *t*_*n*+1_. These rules are detailed in Section A.

**Fig. 4:**
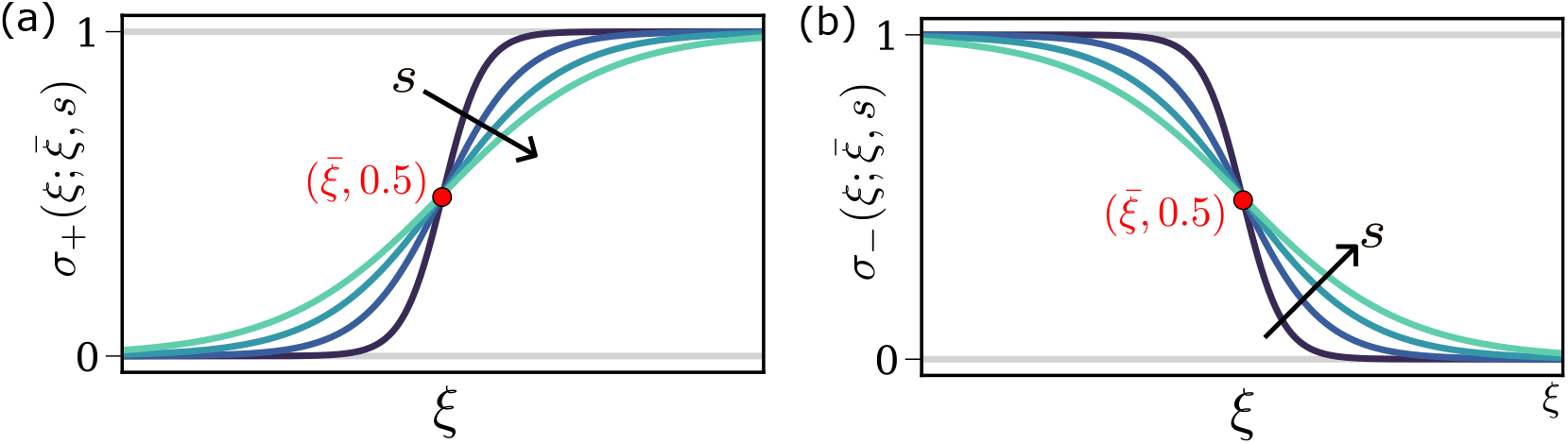
Schematic illustrating how we model the activation (a) and inhibition (b) of cell cycle transition by a variable *ξ*, which can represent both an internal state variable or externally prescribed oxygen levels. The inhibition/activation is modelled using a shifted and rescaled sigmoid function (see Eq. (1)), *σ* parametrized by 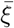, *i.e*., the location of the point of inflation, and *s*, which characterises the steepness of the activation/deactivation curves.

Cells in the *G*_1_ and *C*_1_ states are both in the G1 phase; while *G*_1_ cells have not committed to entering the S phase, *C*_1_ cells have but are transiently arrested due to hypoxia. Transition into, and out of, the *C*_1_ state models hypoxia-mediated activation/deactivation of the G1 checkpoint (see Figure 3). *S* cells remain in this state until they complete DNA replication (*i.e*., when *x* = 2). Cells in the *S* state are sensitive to fork collapse, which occurs when DRF levels drop below a minimal threshold necessary to support the integrity of DNA replication machinery. Cells in states *G*_2_ and *C*_2_ are in the G2/M phase. While cells in *G*_2_ can attempt mitosis, *C*_2_ cells are transiently arrested while they repair accumulated damage (G2 checkpoint). Cell death in the G2/M phase is regulated by a cell damage level, *y*, and can occur either upon transition to the *C*_2_ state or via mitotic catastrophe when *G*_2_ cells that attempt mitosis detect irreparable damage. Upon division, a *G*_2_ cell is replaced by two *G*_1_ cells. All values of their internal variables are inherited from the parent cell, except for the DNA content which is split equally between the two daughter cells (*x* = 1). Dead/senescent cells are instantaneously removed from the population.

### 3.3 Modelling intracellular processes

We account for the impact of hypoxia on DNA synthesis and intracellular damage dynamics by assuming that changes in *m*_dNTP_ and *m*_DRF_ depend on the externally prescribed oxygen levels *c* (see Figure 1). As discussed in Section 2, we assume that expression levels of dNTP and DRF decrease in hypoxia (*c < c*_*H*_), and increase upon reoxygenation (*c > c*_*H*_). Additional noise in the evolution of dNTP and DRF levels is introduced to account for intercellular heterogeneity. Details on the update rules for *m*_dNTP_ and *m*_DRF_ can be found in Section A.3.

#### 3.3.1 Modelling DNA synthesis

The DNA content of cell *i, x*^(*i*)^ ∈ [1, 2], is constant during the G1 (*x*^(*i*)^ = 1) and G2/M (*x*^(*i*)^ = 2) phases. During the S phase, it increases from *x*^(*i*)^ = 1 to *x*^(*i*)^ = 2 at a rate that is assumed to be proportional to its intracellular levels of dNTPs (*i.e*, 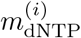). We use the following rule to update the DNA content of cell *i* between times *t*_*n*_ and *t*_*n*+1_:

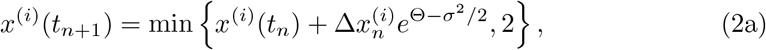

where Θ ∼ 𝒩(0, *σ*) and

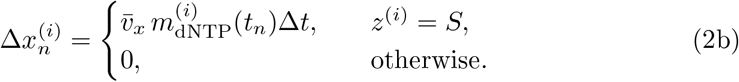

In Eq. (2) the positive constant 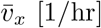 represents the maximum rate of DNA synthesis. The random variable *e*^Θ^ is introduced to capture inter-cellular variability in the rate of DNA synthesis due to factors and mechanisms not captured in the model; the choice of a lognormal noise ensures the physical constraint that DNA can not be degraded (*i.e*., *x*^(*i*)^(*t*_*n*+1_) − *x*^(*i*)^(*t*_*n*_) ≥ 0) and the factor 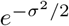 ensures the noise has mean 1. In Figure 5, we show simulations of the DNA dynamics in S phase under different oxygen environments obtained by coupling Eq. (2) to the dynamics of *m*_dNTP_, Eq. (A3), and oxygen levels, Eq. (4).

**Fig. 5:**
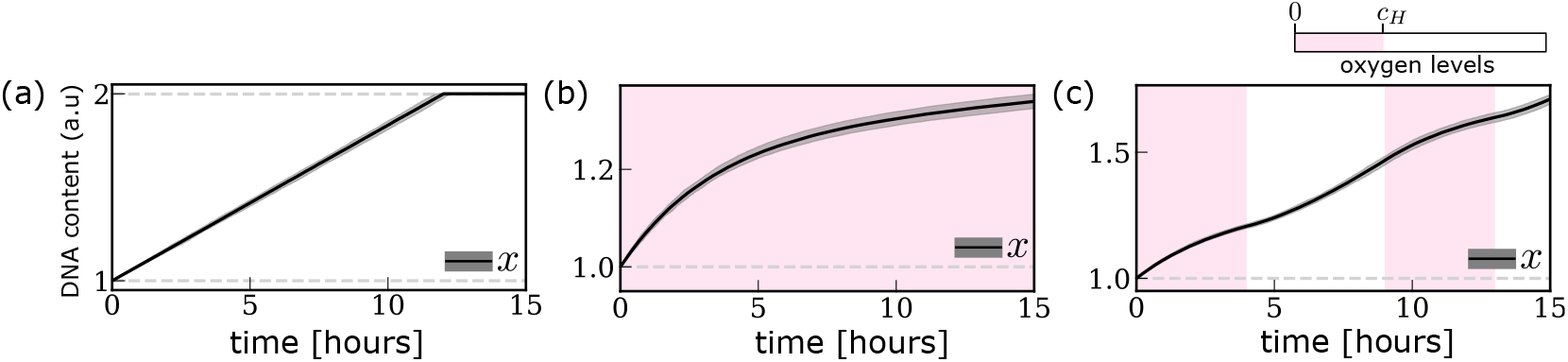
Evolution of DNA content (*x*(*t*)) in the *S* phase under three different oxygen environments: (a) oxygen-rich environment; (b) chronic hypoxia; (c) (4,5)-cyclic hypoxia. The dark line and the shaded grey area indicate respectively the median and 99%-confidence interval for *x*(*t*), measured in arbitrary units (a.u.), and where obtained by simulating Eqs. (2) with *z*^(*i*)^ = *S*, (4) and (A3). The background colours indicate oxygen levels. Parameter values are as indicated in Tables 4-5.

#### 3.3.2 Modelling damage dynamics

We assume that the damage level *y*^(*i*)^ of cell *i* increases as a result of replication stress/damage experienced during the S phase (see Section 2). Damage is repaired during the S phase and via activation of checkpoint signalling (captured in the model via cells transitioning into the *C*_2_ state) in the G2 phase. We assume that in both the S and G2 phases damage repair depends on internal levels of damage repair factors (*m*_DRF_) and that the change in the damage level of cell *i* within a time step Δ*t* satisfies:

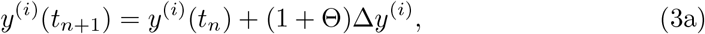

where Θ ∼ 𝒩(0, *σ*) and

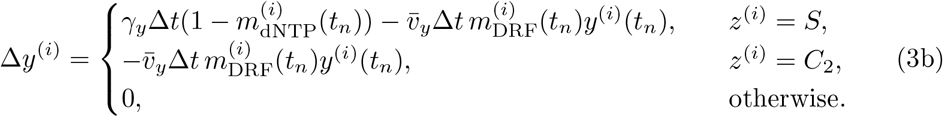

In Eq. (3b), the positive constants 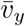 and *γ*_*y*_ represent, respectively, the maximum rate at which damage can be repaired and the rate at which cells accumulate damage due to replication stress. As above, we use multiplicative noise to account for intercellular variability in the damage dynamics (see Eq. (3a)). In writing Eq. (3b), we assume that replication stress is proportional to the slowdown in the rate of DNA replication caused by the drop in intracellular dNTP levels (*i.e*., replication stress 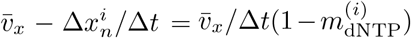). Figure 6 shows simulation of how damage levels in S phase evolve under different oxygen environments. Results are obtained by coupling Eq. (3b) to the dynamics of *m*_dNTP_ and *m*_DRF_ (see Eqs. (A3)-(A4)), and oxygen levels (see Eq. (4)).

**Fig. 6:**
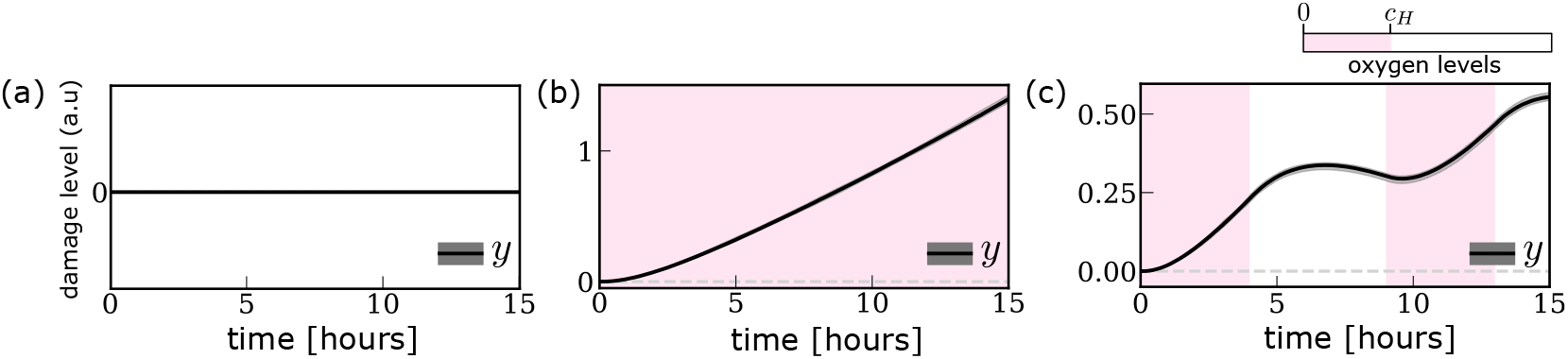
Evolution of damage levels (*y*(*t*)) in three different oxygen environments: (a) oxygen-rich environment; (b) chronic hypoxia; (c) (4,5)-cyclic hypoxia. The dark line and the shaded grey area indicate respectively the median and 99%-confidence interval for *y*(*t*), measured in arbitrary units (a.u.), and were obtained by simulating Eqs. (3b) with *z*^(*i*)^ = *S*, (4) and (A3)-(A4). The background colours indicate oxygen levels. Parameter values are as indicated in Tables 4-5.

### 3.4 Simulation results

Numerical simulations of the IB model are performed in Python. More details on the implementation are given in Appendix A; a pseudocode describing how cell proliferation and death, and intracellular processes are simulated is presented in Algorithms 1 and 2.

We use our IB model to simulate the *in vitro* growth of a population of cancer cells in three oxygen environments:

- **oxygen-rich:**

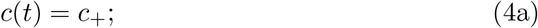
- **chronic hypoxia:**

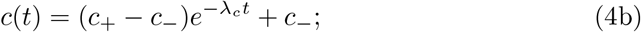
- **cyclic hypoxia:**

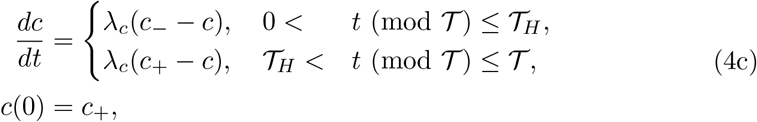

where *t >* 0, the constants *c*_*±*_ (*c*_*−*_ *< c*_*H*_ *< c*_+_) are the minimum and maximum oxygen levels to which cells are exposed, and *λ*_*c*_ is the rate at which oxygen levels relax to their equilibrium values. In Eq. (4c), the function mod indicates the modulus operator, 𝒯 [*hr*] is the periodicity of the fluctuations in oxygen levels, and 𝒯_*H*_ [*hr*] indicates the time of exposure to hypoxia during an oxygen cycle. In other words, cells are repeatedly exposed to 𝒯_*H*_ hours of hypoxia followed by 𝒯_*R*_ = 𝒯 − 𝒯_*H*_ hours of reoxygenation (see Figure 8a). In what follows, the range of possible cyclic hypoxia protocols are characterised by the tuple (𝒯_*H*_, 𝒯_*R*_) and the term (𝒯_*H*_, 𝒯_*R*_)– cyclic hypoxia refers to the oxygen protocol described by Eq. (4c).

As it is standard in *in vitro* experiments, we initialise cells in a regime of (asynchronous) balanced exponential growth (Celora et al, 2022; Webb, 1987) (see Appendix B for details) using the procedure outlined in Algorithm 3. This is the equilibrium regime predicted by the model when cells are exposed to oxygen-rich environments (see Section 4.2). Unless otherwise stated, simulations are initialised with *n*_0_ = 100 cells. For each numerical experiment and set of parameters, we perform 100 realisations of the IB model and use the obtained data to extract the statistical metrics illustrated in Figures 7-11.

**Fig. 7:**
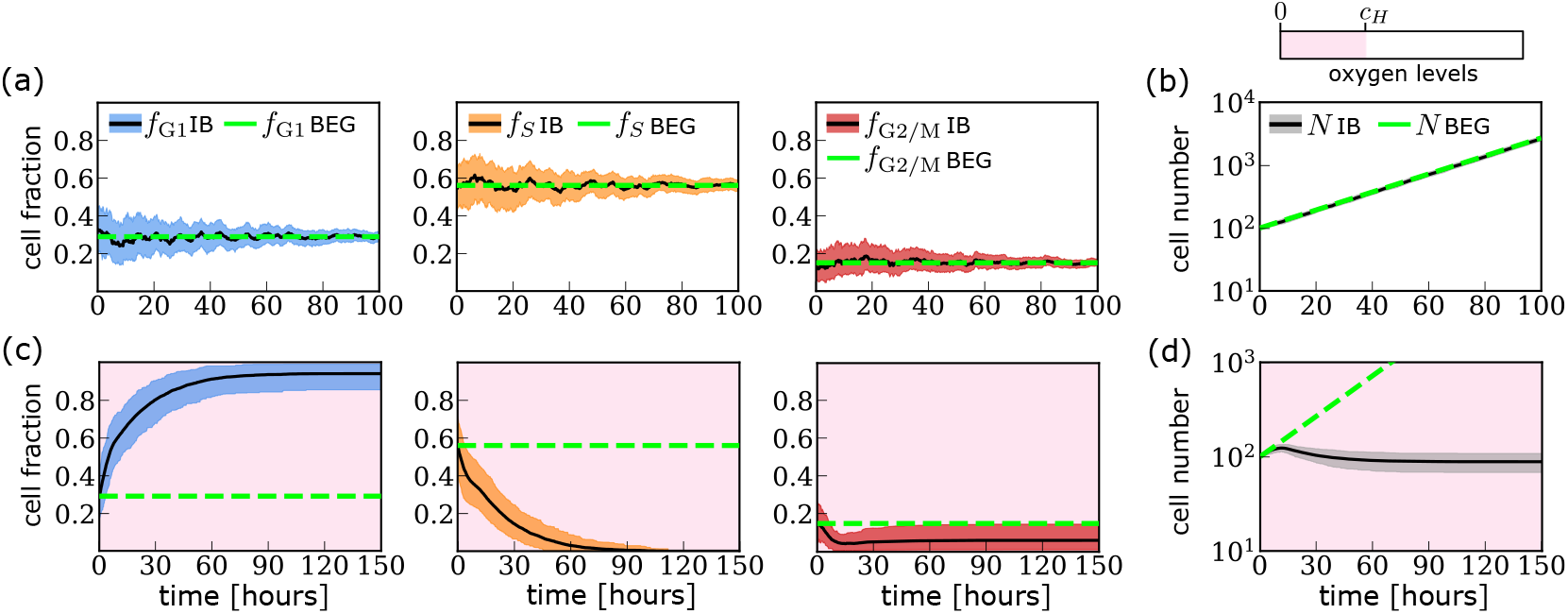
Cell cycle and growth dynamics in constant oxygen environments generated by the IB model. We plot the evolution of the mean and 99%-confidence interval estimates for the evolution of the fraction of cells in each phase of the cell cycle, *f*_*m*_ for *m* ∈ {G1, S, G2/M}, and the total number of cells, *N*, in (a)-(b) oxygen-rich environment and (c)-(d) constant hypoxia. The dashed green lines indicate the analytical prediction from the balanced exponential growth (BEG) model in oxygen-rich environments (see Section B). The background colour indicates oxygen levels. Parameter values for the BEG and IB model are the same and are as indicated in Tables 3-6.

#### 3.4.1 Model parameters

Where possible, model parameters are estimated from the literature, and based on the colorectal RKO cell line which was the focus of previous theoretical (Celora et al, 2022) and experimental studies on cyclic hypoxia (Bader et al, 2021b). See Appendix C for futher details (parameter values given in Tables 3-4).

**Table 3:** Summary of the parameters associated with cell cycle progression, checkpoint dynamics and cell-fate decisions and the values used in the simulations (see Eqs. (A1)-(A2). Typical values (*i.e*., wild-type behaviour) are given for the RKO cancer cell line. Rows highlighted in grey define the parameters that are changed to model populations of cancer cells with different damage repair capacities (see Figure 13). The abbreviation (a.u.) stands for arbitrary units.

|  | Description | Typical value(s) | Justification (Refs) |
| --- | --- | --- | --- |
| $\Delta t$ | Timestep | 1/50 (hours) | sufficiently small to capture the time-scale of all processes |
| $k_1$ | rate at which cells exit $G_1$ | 0.195 (hours$^{-1}$) | (Celora et al, 2022) |
| $K_1$ | maximum rate at which cells leave the G1 checkpoints | 1.00 (hours$^{-1}$) | (Celora et al, 2022) |
| $c_H$ | hypoxia threshold | $\approx 1.0$ (% O$_2$) | (Celora et al, 2022) |
| $s_{C_1}$ | sensitivity of G1 checkpoint to oxygen levels | 0.01 (% O$_2$) | taken to be smaller than $c_H$ |
| $s_{C_2}^{\text{off}}$ | sensitivity deactivation of G2 checkpoint to damage levels | 0.01 (a.u.) | taken to be smaller than $\bar{y}_{C_2}^{\text{off}}$ |
| $K_2$ | maximum rate at which cells leave the G2 checkpoints | 1.00 (hours$^{-1}$) | (taken to be the same as $K_1$) |
| $\bar{y}_{C_2}^{\text{on}}$ | threshold damage level for activation G2 checkpoint | DDR$^- \rightarrow 1.0$ (a.u.)<br>DDR$^{\text{wt}} \rightarrow 0.5$ (a.u.)<br>DDR$^+ \rightarrow 0.1$ (a.u.) | (see Section 3.4.1) |
| $s_{C_2}^{\text{on}}$ | sensitivity of G2 checkpoint activation to cellular damage levels | DDR$^- \rightarrow 0.250$ (a.u.)<br>DDR$^{\text{wt}} \rightarrow 0.125$ (a.u.)<br>DDR$^+ \rightarrow 0.025$ (a.u.) | taken to be $\bar{y}_{C_2}^{\text{on}}/4$ |
| $\bar{y}_{C_2}^{\text{off}}$ | threshold damage level for deactivation of G2 checkpoint | DDR$^- \rightarrow 0.100$ (a.u.)<br>DDR$^{\text{wt}} \rightarrow 0.025$ (a.u.)<br>DDR$^+ \rightarrow 0.005$ (a.u.) | taken to be smaller than $\bar{y}_{C_2}^{\text{on}}$ |
| $k_2$ | rate at which $G_2$-cells attempt mitosis | 0.22 (hours$^{-1}$) | (Celora et al, 2022) |
| $\mu_s$ | maximum rate of cell death due to fork collapse in the S phase | 0.05 (hours$^{-1}$) | match % of apoptotic cells after 20 hr exposure to hypoxia (Bader et al, 2021b) |
| $\bar{m}_{S \rightarrow \emptyset}$ | threshold level of DRFs for cell death in the S phase | 0.22 (a.u.) | corresponds to $m_{\text{DRF}}$ levels after 12 hr in hypoxia (see Figure 14) |
| $\bar{y}_{M \rightarrow \emptyset}$ | threshold damage level for mitotic catastrophe | 1.00 (a.u.) | normalised |
| $\bar{y}_{G2 \rightarrow \emptyset}$ | threshold level damage level for cell death upon G2 checkpoint activation | 2.50 (a.u.) | taken to be greater than $\bar{y}_{M \rightarrow \emptyset}$ |
| $\epsilon_{S \rightarrow \emptyset}$ | sensitivity of S-cell death to changes in DRF levels | 0.05 (a.u.) | taken to be less than half of $\bar{m}_{S \rightarrow \emptyset}$ |
| $\epsilon_{M \rightarrow \emptyset}$ | sensitivity of mitotic catastrophe to damage levels | 0.25 (a.u.) | taken to be less than half of $\bar{y}_{M \rightarrow \emptyset}$ |
| $\epsilon_{G2 \rightarrow \emptyset}$ | sensitivity of cell-death upon G2 checkpoint activation to damage levels | 1.00 (a.u.) | taken to be less than half of $\bar{y}_{G2 \rightarrow \emptyset}$ |

We account for different regulation of damage repair by varying parameters modulating G2 checkpoint activation (*i.e*., probability of cell transitioning into and out of the cell cycle state *C*_2_) in response to damage, see Eq. (A1d); namely, parameters 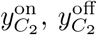 and 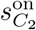 (see Table 3). We model cells with enhanced damage repair activity (DDR^+^ cells) compared to the reference (or wild-type) behaviour (DDR^wt^ cells) by decreasing 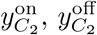 and 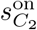 relative to their default values; we account for defective damage repair activity (DDR^-^ cells) compared to wild-type behaviour (DDR^wt^ cells) by increasing 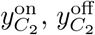 and 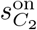 relative to their default values. For more details, see Section C.1.

#### 3.4.2 Clonogenic assays

We estimate cancer cell survival in cyclic hypoxia by developing *in silico* clonogenic assay that is based on a standard “plating before treatment approach” (Franken et al, 2006). This means that cells are first plated and then exposed to cyclic hypoxia. After being exposed to cyclic hypoxia for *t*_*R*_ = 15 (𝒯_*H*_ + 𝒯_*R*_) hours, cells are cultured in ambient oxygen conditions (21%O_2_) for 10 days. The survival fraction is estimated at the end of the 10 days as the ratio

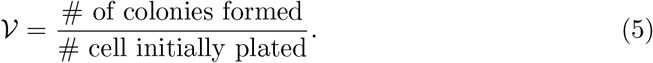

In Eq. (5) we define a colony as a cluster of at least 50 cells that originate from the same progenitor. Note that, in writing Eq. (5), we follow the standard convention by defining survival as the ability of cells to escape replicative death and maintain uncontrolled proliferation when exposed to toxic agents (here cyclic hypoxia). We remark that replicative death can be due to cell death but also persistent cell cycle arrest.

#### 3.4.3 Serial passage assays

We estimate the relative fitness of distinct cancer cell lines under different cyclic hypoxia conditions by simulating serial passage assays. We simulate co-cultures of three cell lines; namely DDR^+^, DDR^-^ and DDR^wt^ (modelled by changing parameter values as described in Section 3.4.1). We initialise the model with 150 cells from each cell line, for a total of 450 cells. While being exposed to a specific cyclic hypoxia protocol, cells are “passaged” every ⌊48*/*𝒯 ⌋ 𝒯 (*i.e*., approximately every 2 days) where ⌊ · ⌋ indicates the floor function. Replating is simulated by randomly sampling *n*_0_ cells from the population; oversampling is used, if the size of the population at the time of the replating is less than *n*_0_. This procedure artificially introduces a carrying capacity and enhances the positive selection pressure on cells that are more adapted to cyclic hypoxia. The fitness of DDR^*±*^ cells relative to DDR^wt^ is quantified by estimating the ratio

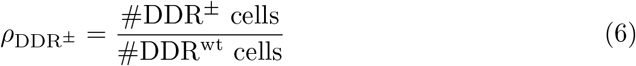

after passaging the population 10 times. DDR^*±*^ cells have a fitness advantage in cyclic hypoxia over DDR^wt^ cells if *ρ*_DDR_*± >* 1 (and conversely). Since *ρ*_DDR_*±* are stochastic variables, statistical evidence for the alternative hypothesis *ρ*_DDR_*±* ≤ 1 and *ρ*_DDR_*±* ≥ 1 is tested using one-sample one-tail t-test with p-value 0.001.

## 4 Results

We use our IB model to simulate cancer cell responses to different oxygen environments. In Section 4.1, we demonstrate that the IB model reproduces the cell cycle and population growth dynamics observed *in vitro* under constant high-oxygen and constant hypoxic conditions. In Section 4.2, we simulate cell culture and clonogenic assay experiments in a wide range of cyclic hypoxia environments. We identify a range of population-level dynamics for different cyclic hypoxia protocols: sustained growth, dormancy and population extinction. Using the IB model, we can relate population-level behaviour to the dynamics of individual cell states and specifically their damage regulation. Finally, in Section 4.3, we study how damage repair capacity influences cancer cell fitness under different cyclic hypoxia conditions by simulating serial passage assays.

### 4.1 Model predictions in constant environments

We validate our IB model by simulating cell cycle and growth dynamics of a population of wild-type cancer cells under constant oxygen conditions. The results are shown in Figure 7.

As expected, in the oxygen-rich environment (see Figures 7a-7b) the model predicts balanced exponential growth (note that in Figure 7b we use a log scale for the y-axis). The total number of cells *N* eventually grows exponentially at a constant rate *λ*_BEG_, while the fraction of cells in each cell cycle phase asymptotes to an equilibrium value, 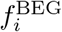 for *i* ∈ {G1, S, G2/M}, with uncertainty in the values of the cell fractions *f*_*i*_ decreasing over time. This is in line with the predictions of the deterministic model in (Celora et al, 2022) (the relationship between the two models is discussed in Appendix B).

Under constant hypoxia (see Figures 7c-7d), after an initial transient, the average number of cells evolves to a constant value. While the number of cells increases for the first ≈ 12 hours, it then decreases due to the death of cells in the S phase under chronic hypoxia. At long times, most cells are in the G1 phase having successfully arrested in the G1 checkpoint, (*i.e*., they are locked in state *C*_1_). The delayed decrease in population size due to the death of cells in the S phase and the predicted accumulation of quiescent cells agree with constant hypoxia experiments on RKO cell line culture (Bader et al, 2021a).

### 4.2 Characterising the wild-type responses to different cyclic hypoxia environments

#### 4.2.1 Growth dynamics

We use the IB model to simulate the cell cycle and growth dynamics of a population of cancer cells under a range of cyclic hypoxia conditions. The results presented in Figure 8 show that the long-term growth dynamics depend on the oxygen protocol used. We characterise these dynamics by estimating the asymptotic population growth rate, *λ* (see Figure 8b)). This is defined by fitting an exponential function to the change in population size over a period 𝒯 (see schematic in Figure 8a). Asymptotically, and assuming a sufficiently large population of cells, the estimated *λ* is expected to be independent of the time *t* chosen for its estimation.

**Fig. 8:**
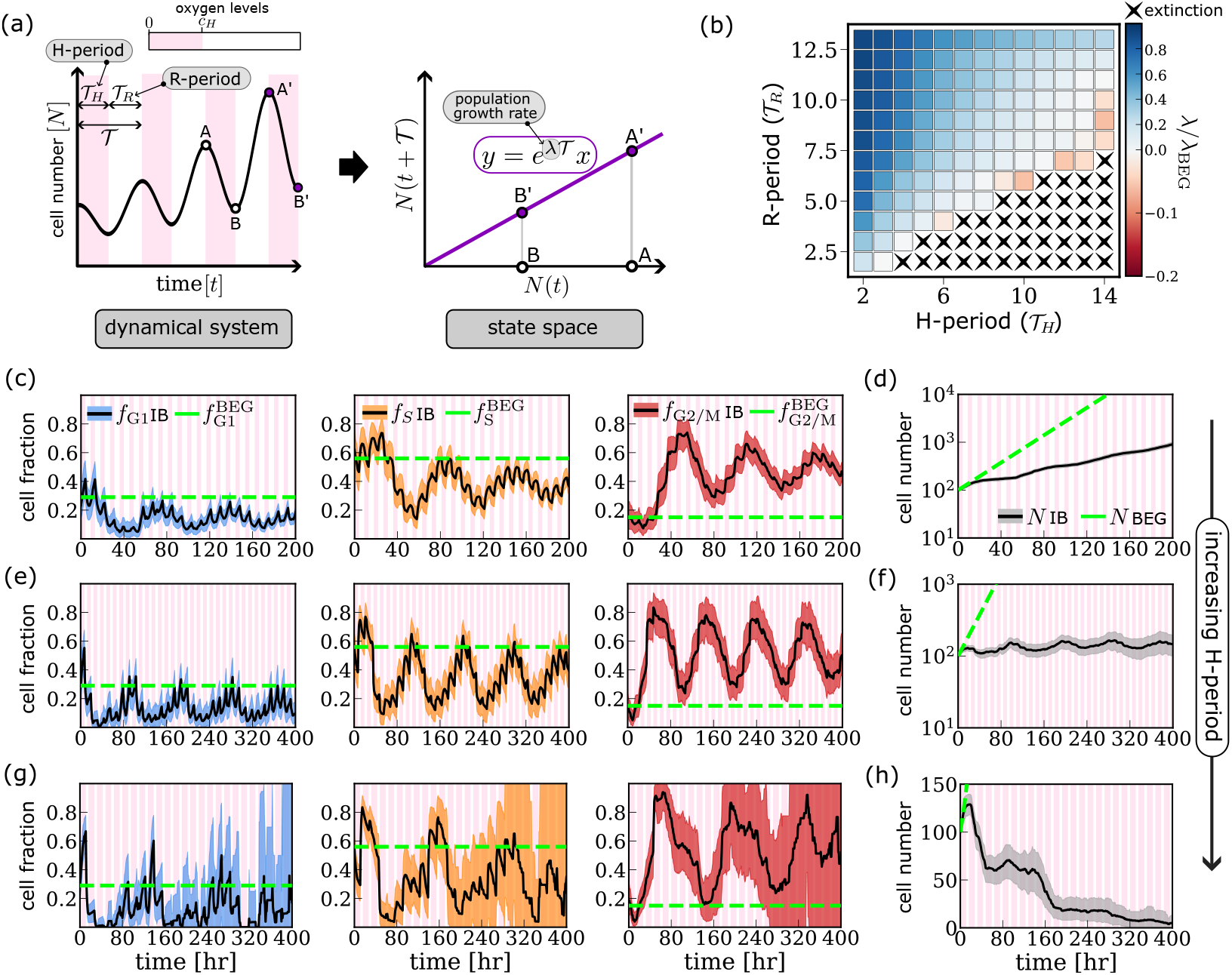
Cell cycle and tumour growth dynamics *in vitro* under different cyclic oxygen environments. (a) Schematic showing how the population growth rate is defined in a fluctuating environment (*i.e*., *dN/dt* = *r*(*t*)*N* where *r* is 𝒯 -periodic function). The time-evolution of cell number *N* deviates from the exponential growth model (see left-hand side plot). However, when projected onto the (*N* (*t*),*N* (*t* + 𝒯)) state space, the behaviour is analogous to one of exponential growth population in a constant environment (see right-hand side plot). (b) Estimated population growth rate *λ* for a range of cyclic hypoxia protocols. Crosses indicate conditions for which the cell population goes extinct with probability ≥ 90%. (c)-(h) We plot the evolution of the fraction of cells in each phase of the cell cycle, *f*_*m*_ with *m* ∈ {G1, S, G2/M}, and the total number of cells, *N*, as predicted by the IB model for (c)-(d) (4,5)–cyclic hypoxia; (e)-(f) (7,5)–cyclic hypoxia; and (g)-(h) (11,5)–cyclic hypoxia. Parameter values, variables and colours are as in Figure 7.

When 𝒯_*H*_ is sufficiently short, the model predicts sustained population growth, albeit at a lower rate than in oxygen-rich conditions (*i.e*., during the balanced exponential growth regime), *i.e*., 0 *< λ* ⪅ *λ*_BEG_; this is the case, for example, when cells are exposed to (4,5)–cyclic hypoxia (see Figure 8d). When considering the corresponding cell cycle dynamics, despite the persistent fluctuations in the cell cycle fractions, we observe a systematic increase in the fraction of cells in the G2/M phase, *f*_G2/M_ (see Figure 8f). We note that the period of the fluctuations in *f*_G2/M_ is eventually the same as for the oxygen levels – in this case 9 hours. As 𝒯_*H*_ increases, the IB model predicts substantial inhibition of population growth (or growth arrest); this is the case, for example, when simulating exposure of cells to (7,5)–cyclic hypoxia (see Figure 8f). Recall that under constant hypoxia, growth inhibition is due to cells arresting in the G1 phase. By contrast, under (7,5)-cyclic hypoxia, cells continue proceeding through the cell cycle (compare Figures 7d and 8e) suggesting that cell proliferation continues even though the total number of cells in the population is not increasing. There are two possible causes of population growth inhibition (or population dormancy) (Wells et al, 2013): cell cycle arrest or a balance between cell death and proliferation. We conclude that population dormancy in (7,5)–cyclic hypoxia is due to an increase in cell death. Finally, when considering cyclic hypoxia protocols with larger 𝒯_*H*_, the model predicts population extinction with high likelihood (see crosses in Figure 8b); this is the case, for example, when simulating exposure to (11,5)–cyclic hypoxia (see Figure 8h). We note that the uncertainty in model predictions for the cell cycle dynamics increases on the long-time scale. The widening of the confidence intervals in Figure 8g is due to the increased dominance of demographic noise as the number of cells approaches zero. Nonetheless, all replicates eventually predict population extinction.

#### 4.2.2 Cell survival

In the simulations presented in Section 4.2.1, we investigated the impact of fluctuating oxygen levels on the emergent population growth dynamics. In this section, we investigate cell survival in different oxygen environments and its relation to observed growth dynamics and cell survival by simulating clonogenic assay experiments (see Section 3.4.2).

In Figure 9a, we report estimated values of the survival fraction 𝒱 for different cyclic hypoxia protocols. As in Figure 8, we characterise cyclic hypoxia environments by the hypoxia period, 𝒯_*H*_, and the reoxygenation period, 𝒯_*R*_. We find significant variation in the survival fraction significantly as 𝒯_*H*_ and 𝒯_*H*_ vary. For sufficiently large reoxygenation periods 𝒯_*R*_, cells are likely to survive and 𝒱 ≈ 1. In contrast, cells are more likely to die than survive when the reoxygenation period is short and the hypoxia period is sufficiently long. Overall, our results highlight that both the overall time of exposure to hypoxia and the evolution of the oxygen dynamics are important in determining the extent to which hypoxia is toxic for cells.

**Fig. 9:**
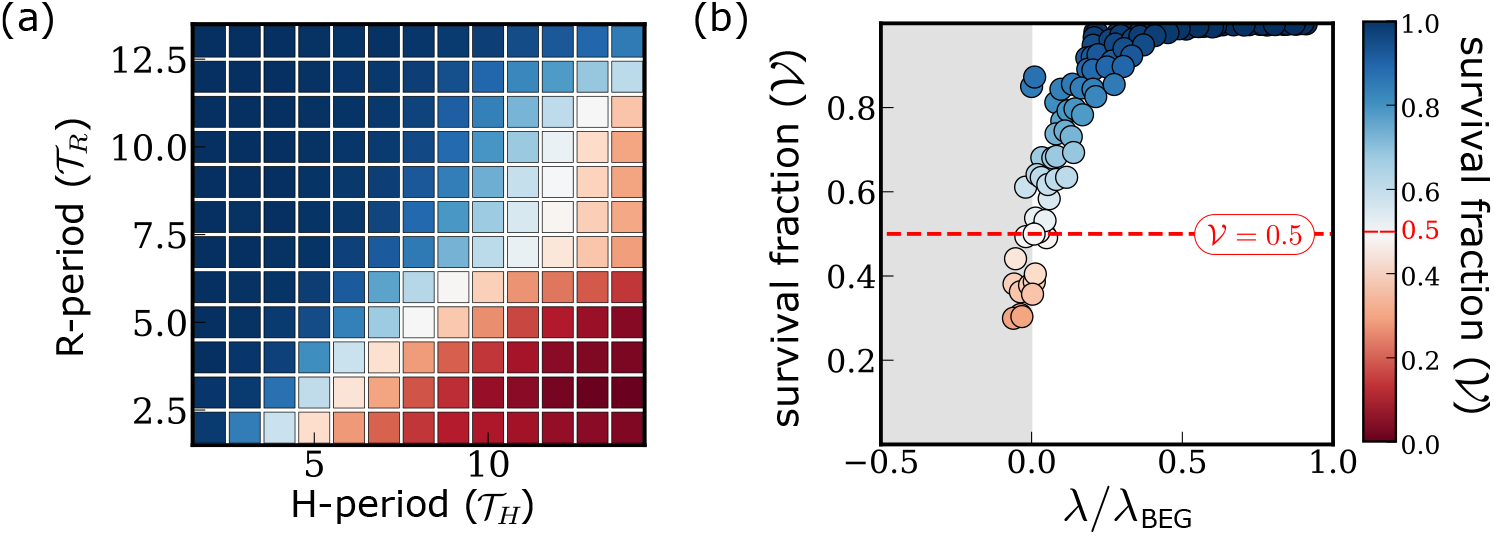
Characterising wild-type cell survival in different oxygen environments. (a) Mean estimator for the cell survival fraction 𝒱 (see Section 3.4.2) in a range of cyclic hypoxia conditions. (b) Scatter plot illustrating the relation between the population growth rate *λ* (see Figure 8b) and cell survival for the cyclic hypoxia conditions studied in (a) except those that lead to extinction of the cell population (see Figure 8b). Parameters are as indicated in Tables 3-6.

The results presented in Figure 9b suggest that estimates of cell survival and population growth rates in cyclic hypoxia are related. In all conditions where we predict a positive growth rate, the survival fraction never drops below 0.5 (see red line in Figure 9b). This suggests that must be more likely to survive than to die to avoid population decay or extinction. While this is intuitive when considering a homogeneous population in which the survival probability is the same for all cells, in our model, cell cycle heterogeneity influences a cell’s survival probability (see Appendix D). This correlation is lost when cells are exposed to fluctuating oxygen levels for sufficiently long times. The decay time scale for such correlations depends on the oxygen dynamics and tends to infinity when 𝒯_*R*_ → 0 (*i.e*., under chronic hypoxia). This is because, in our model, the G1 checkpoint arrest is irreversible (see Figure 7c) so that the estimates of cell survival are determined, even at long times, by the initial cell cycle distribution.

Overall, a decrease in cell survival 𝒱 corresponds to a decrease in the population growth rate *λ*. Nonetheless, we identify a significant range of environmental conditions in which the population growth rate *λ* decreases even though 𝒱 ≈ 1. In these cases, the reduction in the cell proliferation rate is driven by the activation of cell cycle checkpoints and the consequent increase in cell cycle duration (see, for example, Figure 7c-7d).

### 4.3 Characterising the link between damage repair capacity and cancer cell responses to cyclic hypoxia

In the previous section, we showed how the response of a cancer cell line to cyclic hypoxia depends on how the oxygen levels fluctuate (*i.e*., the values of 𝒯_*H*_ and 𝒯_*R*_). Based on these results, we now partition the (𝒯_*H*_, 𝒯_*R*_) parameter space into four regions depending on the predicted cell responses (see Figure 10a). As 𝒯_*R*_ decreases and 𝒯_*H*_ increases (*i.e*., transitioning from the dark green to the dark pink regions in Figure 10a), the environmental conditions become increasingly toxic for cancer cells. Genetic and phenotypic heterogeneity in the regulation of DNA damage response (DDR) and cell-cycle checkpoints signalling has been observed in solid tumours (Begg and Tavassoli, 2020; Jiang et al, 2020). This includes: alterations that silence DDR signalling (Jiang et al, 2020) (DDR^-^ cells), thereby allowing cells to proliferate faster by suppressing damage repair signalling; and, alterations that enhance DDR signalling (Wu et al, 2023) (DDR^+^ cells), thereby promoting cell repair signalling and survival, and, therefore, resistance to chemo- and radiotherapy. We simulate serial passage experiments to investigate how these alterations to damage repair signalling affect cancer cell fitness in different fluctuating oxygen environments (see Section 3.4.3).

**Fig. 10:**
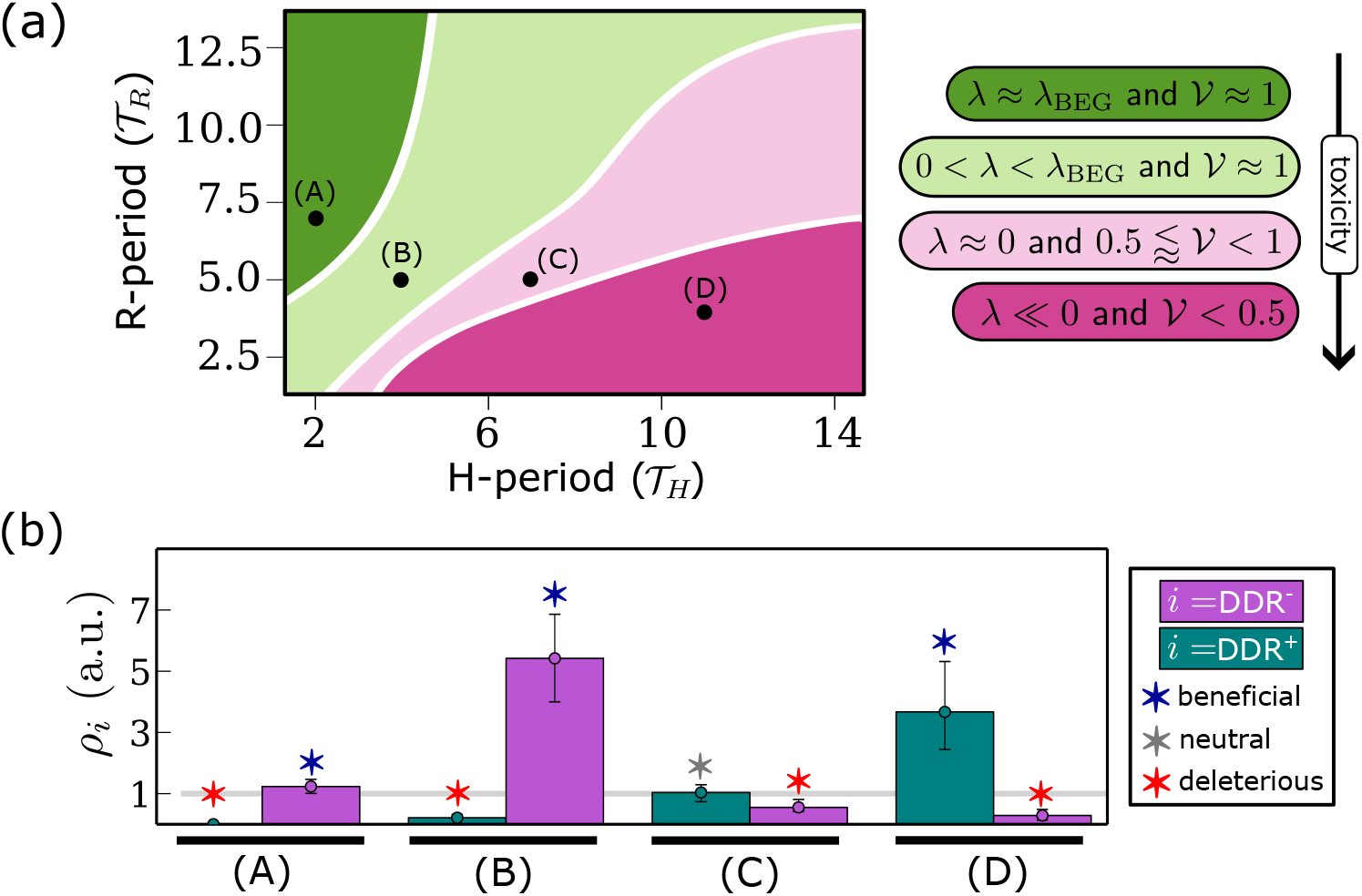
(a) Schematic summarising the characteristic wild-type responses to different cyclic hypoxia protocols. We decompose the (𝒯_*H*_, 𝒯_*R*_) space into four regions characterised by the estimated values of the population growth rate *λ* (see Figure 8) and survival fraction 𝒱 (see Figure 9). (b) Histograms showing the relative fitness *ρ*_DDR_*±* (see Eq. (6)) for different cyclic hypoxia protocols: (A) (2,7)–cyclic hypoxia; (B) (4,5)–cyclic hypoxia; (C) (7,5)–cyclic hypoxia; (D) (11,4)–cyclic hypoxia. Red, grey and blue stars indicate DDR alterations that are, respectively, beneficial (*ρ*_*i*_ *>* 1, p-value=0.001), deleterious (*ρ*_*i*_ *<* 1, p-value=0.001) or neutral (if neither beneficial or deleterious). More details on how we quantify relative fitness are given in Section 3.4.3. Parameter values are as indicated in Tables 3-6.

The results are presented in Figure 10b. Overall, we find that the estimated relative fitness *ρ* of both DDR^+^ and DDR^-^ cells depends on the cyclic hypoxia protocols. For DDR^+^ cells, *ρ* increases as the extent to which cyclic hypoxia is toxic for wild-type cells increases. In contrast, *ρ*_DDR_*−* depends non-monotonically on cyclic hypoxia toxicity, being maximal for midly-toxic conditions and 𝒯_*H*_ sufficiently small (*i.e*., condition (A) in Figure 10b).

Under cyclic hypoxia conditions that are harmless for wild-type cells (dark green region in Figure 10a; condition (A) in Figure 10b), enhanced damage repair capacity is deleterious (*ρ*_DDR_+ *<* 1), while deficiencies in damage repair capacity are mildly beneficial (*ρ*_DDR_*−* ⪆ 1). In contrast, under cyclic hypoxia conditions that are highly toxic for wild-type cells (dark pink region in Figure 10a; condition (D) in Figure 10b), enhanced activation of the DDR increases cell fitness (*ρ*_DDR_+ *>* 1), while deficiencies in the DDR are significantly deleterious for cells. Between these two extremes (*i.e*., protocols within the light green and light pink regions of the schematic in Figure 10a; condition (B)-(C) in Figure 10b), the model predicts that enhanced damage repair capacity switches from being deleterious to being beneficial, while deleterious damage repair capacity becomes increasingly deleterious. Interestingly, these transitions are characterised by regimes in which the composition of the population at the end of the simulations is highly heterogeneous, with coexistence of DDR^+^, DDR^*−*^ and DDR^wt^ cells even after several passages. For example, in case (C) in Figure 10b, the coexistence of different cell types is reflected in *ρ*_DDR_+ and *ρ*_DDR_*−* being both close to one. This suggests that cyclic hypoxia can give rise to conditions in which the fitness landscape associated with DDR regulation is flat and, consequently, natural selection is very slow.

#### 4.3.1 Heterogeneity in cell damage repair capacity shapes damage distribution under cyclic hypoxia

To better understand the relation between damage repair and cell fitness in cyclic hypoxia, we look at how selection reshapes the damage distribution within the cell population. We quantify this by comparing the cell damage distribution in the coculture experiment with the distribution under control conditions where serial passage assays are only performed with DDR^wt^ cells (*i.e*., default values of the model parameters). The use of the control case for comparison is important since “passaging” of cells can alter both the cell cycle and cell damage distributions. In general, because cells are exposed to fluctuating oxygen levels, the damage distribution eventually converges to a periodic function that fluctuates with the same frequency as the passaging, (see Section E). The results, therefore, depend on the time *t* at which the damage distribution is computed. Here we focus on the damage distribution at the end of the final hypoxic phase:

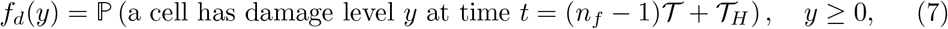

where *n*_*f*_ = 10⌊48*/*𝒯⌋ indicates the total number of oxygen cycles to which the cells have been exposed during the serial passage assay. This is the time at which selection (via passing) operates.

The results presented in Figure 11 show that damage levels in the co-culture and control experiments can differ markedly depending on the cyclic hypoxia protocol considered. For (2,7)-cyclic hypoxia, despite the reported mild advantage of DDR^-^ cells (see (A) Figures 10b), damage levels are comparable between the co-culture and control experiments. The brief exposure to hypoxia is not sufficient to drive significant damage accumulation even in cancer cells with deficiencies in damage repair capacity. When considering (4,5)-cyclic hypoxia, DDR^-^ cells are predicted to have a significant advantage (see (B) Figure 10b). However, unlike in (2,7)–cyclic hypoxia, accumulation of DDR^-^ cells is associated with the median damage level in co-culture conditions being significantly higher than in the control conditions. In this intermediate regime, deficiencies in damage repair capacity are most beneficial for cancer cells as they promote high proliferation but at the cost of cells accumulating significantly higher damage/stress. For more toxic cyclic hypoxia conditions (*i.e*., light and dark pink region in Figure 10a), the trend reverses; lower levels of damage are recorded for the co-culture than control conditions (see results for (C) (7,5)-cyclic hypoxia and (D) (11,4)-cyclic hypoxia in C11). The environmental toxicity is such that there is no benefit in attempting to proliferate; and cells are better off prioritising their survival by enhancing damage repair signalling and, thereby, maintaining low damage levels (see Figure 10).

**Fig. 11:**
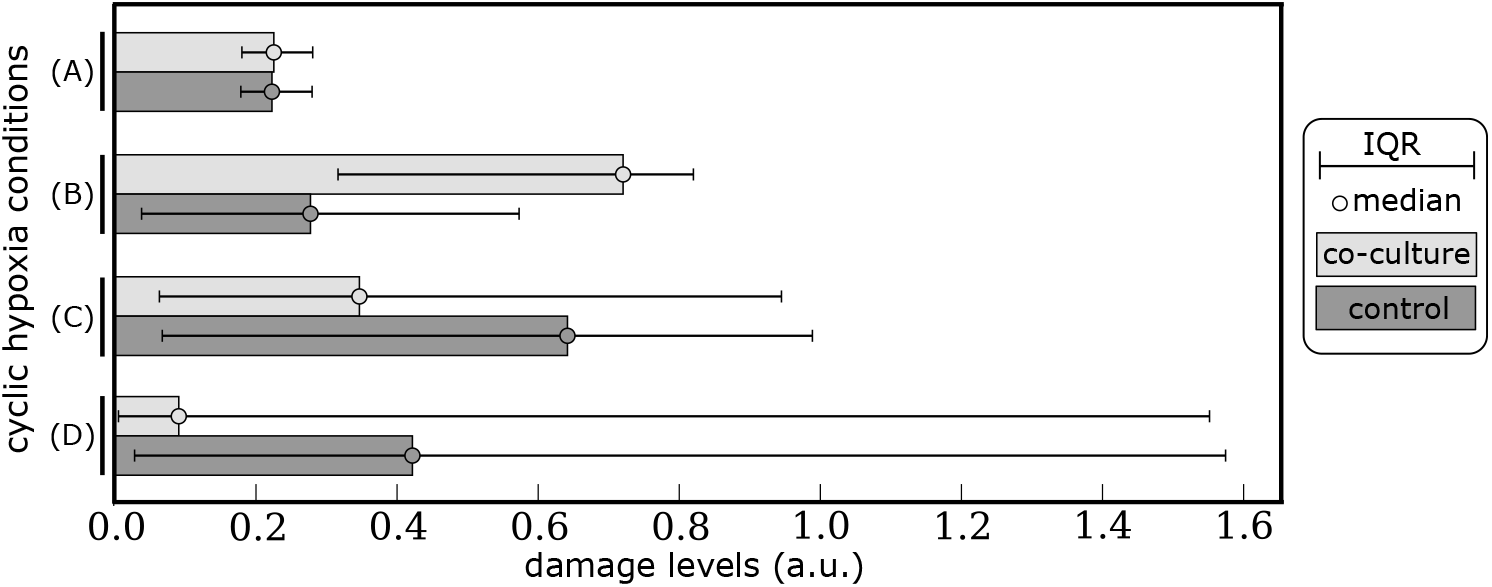
Median *m*_*D*_ and interquartile range IQR_*D*_ of the damage distribution (defined in Eq. (7)) for co-culture (light grey) and control (dark grey) conditions for cells grown in the same cyclic hypoxia conditions investigated in Figure 10: (A) (2,7)–cyclic hypoxia; (B) (4,5)–cyclic hypoxia; (C) (7,5)–cyclic hypoxia; (D) (11,4)–cyclic hypoxia. Parameters are as indicated in Tables 3-6.

## 5 Application of the result to intra-tumour heterogeneity

Changes to the regulation of cell cycle checkpoints and DNA repair pathways are common in cancer as they sustain uncontrolled proliferation (Viner-Breuer et al, 2019; Jiang et al, 2020). However, in conditions where proliferation can not be sustained, functioning checkpoint regulation can play an essential role in favouring cancer cell survival. For example, in tumour regions that are chronically exposed to severe hypoxia, cells experience replication stress and, therefore, activation of cell cycle checkpoints in response to low oxygen levels (*i.e*., hypoxic stress) can be crucial for cancer cell survival (Qiu et al, 2017; Pires et al, 2010b,a). The different roles of cell cycle checkpoints in well-oxygenated and hypoxic regions influence the regulation of cancer cell proliferation, thus favouring heterogeneity in hypoxic tumours (Begg and Tavassoli, 2020; Emami Nejad et al, 2021).

Heterogenous blood flow in vascularised tumours can generate tissues in which oxygen levels fluctuate, exposing cells to cyclic hypoxia. The period and amplitude of such fluctuations may vary with the distance from the closest vessel. In (Ardaševa et al, 2020), it was proposed that spatiotemporal variability in oxygen levels creates ecological niches that foster intratumour phenotypic heterogeneity along the cell metabolic axis. Our results suggest that spatiotemporal heterogeneity in tumour oxygen levels can contribute to intratumour heterogeneity in damage repair capacity. This is because the impact of damage repair capacity on cellular fitness under cyclic hypoxia depends on both the frequency and duration of hypoxia periods (see Figure 10). While deficient damage repair capacity is advantageous for cancer cells when hypoxia periods are rare, it is deleterious when hypoxia periods are long and frequent. Under such environmental conditions, enhanced damage repair capacity is necessary to sustain prolonged checkpoint activation and allow cell survival. Interstingly, we can identify intermediate cyclic hypoxia conditions under which cancer cells with different damage repair capabilities coexist. This suggests that cells with enhanced damage repair capacity, which is usually associated with resistance to treatment, may be located in regions that are primarily hypoxic and highly toxic for cancer cells, and also in regions that are frequently reoxygenated and can sustain cancer cell proliferation albeit at a lower rate than better oxygenated areas.

## 6 Conclusion

We have developed a stochastic, individual-based (IB) model of *in vitro* cancer cell cultures to study the impact of hypoxia-driven cell cycle dysregulation on cancer cell responses (*e.g*., proliferation, survival and damage regulation) to different dynamic oxygen environments. Our model extends previous work on modelling cell cycle progression in cyclic hypoxia (Celora et al, 2022) by coupling cell cycle progression to damage repair dynamics. Interestingly, we find that cancer cell responses to dynamic oxygen environments dynamics of the oxygen levels and that fluctuating oxygen environments can drive variability in cell damage repair capacity within tumours.

Our model describes how oxygen fluctuations impact cell cycle progression and cell survival by affecting DNA replication and repair within cells. In Section 4.2, we showed the cell cycle and growth dynamics predicted by the model for different oxygen environments. In constant oxygen environments, the model reproduces the expected population dynamics for non-confluent cell cultures: balanced exponential growth and growth arrest driven by cell quiescence in the G1 phase. Interestingly, we found that, depending on the duration and frequency of hypoxia periods, cyclic hypoxia can yield very different growth patterns in the same cancer cell population: exponential growth, saturated growth or population extinction. By simulating *in vitro* clonogenic assays, we were further able to characterise cell survival in different cyclic hypoxia conditions. By combining population growth rate and survival estimates, we partitioned the space of possible cyclic hypoxia protocols (*i.e*., (𝒯_*H*_, 𝒯_*R*_)-space) into four regions each associated with qualitatively distinct cancer cell responses (see Figure 10a).

In Section 4.3, we studied how alterations to DNA damage response and cell cycle checkpoint signalling (or, in brief, cell damage repair capacity) influence cancer cell responses to cyclic hypoxia. We considered two types of alterations: deficient damage repair capacity (DDR^-^), which promotes uncontrolled proliferation while hindering damage repair; and enhanced damage repair capacity (DDR^+^), which promotes damage repair and cell survival. Our results suggest that cyclic hypoxia may define different environmental niches: those in which either DDR^+^ or DDR^-^ cells localise; and those in which cells with different DDR signalling coexist. We concluded by discussing the predictions of our model in the context of intratumour heterogeneity in vascularised tumours.

There are several ways in which our work could be extended. We developed our IB model to replicate *in vitro* monolayer conditions, to allow for comparison and validation with experimental data. While cell culture experiments are effective for building a mechanistic understanding, they can not capture the complexity of interactions and 3D organisation of tumours growing *in vivo*. Hence, there is a growing interest in advancing 3D tumour cultures, such as multicellular spheroids and organoids, to bridge the gap between *in vitro* and *in vivo* conditions. A natural extension of our work would be to integrate our cell cycle model within a multiscale framework to study spheroid growth, using either IB (Bull et al, 2020; Hamis et al, 2021; Ghaffarizadeh et al, 2018; Jiménez-Sánchez et al, 2021) or continuous modelling (Murphy et al, 2023; Pérez-Aliacar et al, 2023) approaches.

The results from the *in silico* serial passage assays highlight the role of damage repair capacity in shaping cancer cell responses and adaptation in fluctuating oxygen environments. Here, we have modelled pre-existing alteration of the damage repair capacity of cells, neglecting behavioural changes that may occur during the time scale of the experiments. In practice, cell cycle progression and damage repair are regulated at the genetic and epigenetic levels. While genetic mutations lead to irreversible changes in cancer cell behaviour, epigenetic mutations are reversible and dynamically regulated. Building on previous work on structured-population modelling (Ardaševa et al, 2020; Celora et al, 2023; Lorenzi and Painter, 2022), it would be interesting to investigate the interplay between damage repair, phenotypic heterogeneity and cyclic hypoxia in shaping intratumour heterogeneity.

## Supplementary information

The article has no accompanying supplementary files

## Acknowledgments

GLC is supported by funding from the Engineering and Physical Sciences Research Council (EPSRC) and the Medical Research Council (MRC) (Grant No. EP/L016044/1 and Grant No. EP/W524335/1). RN is supported by the Engineering and Physical Sciences Research Council (Grant No. EP/W524311/1).

## Appendix A Detailed implementation of *in vitro* cancer cell dynamics in hypoxi

We detail the implementation of cell proliferation and death, and intracellular processes in our individual-based (IB) model in Algorithm 1 and Algorithm 2. As shown in Figure 3, for each cell, we first update its cell cycle state (as discussed in Section A.1). We then check for cell division/death (as discussed in Section A.2) and finally update its internal variables (as discussed in Section A.3). Given a sufficiently small time step Δ*t* and denoting by *c*_*n*_ the oxygen levels at time *t*_*n*_, the following rules are used to simulate cell cycle progression and cell fate decisions (*i.e*., cell death) in the time interval [*t*_*n*_, *t*_*n*_ + Δ*t*):

### A.1 Cell-cycle transitions and checkpoint dynamics

At each time step, any surviving cell in *G*_1_, *C*_1_, *S*, and *C*_2_ can transition to the next cell cycle phase, arrest due to activation of a checkpoint or re-enter the cell cycle upon checkpoint deactivation. The dynamics of cells in states *G*_1_ and *C*_1_ cells are modelled as in (Celora et al, 2022). New rules are introduced to describe the evolution of *S, G*_2_ and *C*_2_ cells (see Section A). Based on experimental evidence (see Section 2), cell cycle progression is implemented as follow

- A cell in state *z*^(*i*)^ = *C*_1_ may re-enter the cell cycle by transitioning to the S phase with probability

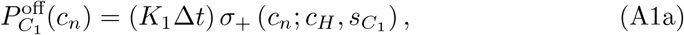

where *σ*_+_ is given by Eq. (1), *c*_*H*_ is the threshold oxygen levels for hypoxia, and the positive constant *K*_1_ denotes the maximum rate at which cells exit the *C*_1_ state and initiate DNA synthesis by entering the *S* state. Eq. (A1a) captures the inhibitory effect of hypoxia (*c < c*_*H*_) on the *C*_1_ → *S* transition; based on previous work (Celora et al, 2022), we assume a switch-like behaviour and fix 0 *< s*_*C*_1 ≪ 1.
- A cell in state *z*^(*i*)^ = *G*_1_ may arrest in the G1 phase (*z*^(*i*)^ → *C*_1_) or proceed to the S phase (*z*^(*i*)^ → *S*) with probabilities

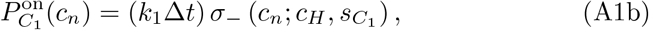

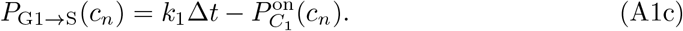 In Eqs. (A1b)-(A1c), the positive constant *k*_1_ represents respectively the rate at which cells exit the G1 phase in oxygen-rich conditions, while *c*_*H*_ and *s*_*C*_1 are as above. In Eq. (A1b), the hypoxia-mediated activation of the G1 checkpoint is captured by the *σ*_*−*_ activation function, which is defined as in Eq. (1).
- A cell in state *z*^(*i*)^ = *C*_2_ may exit G2 arrest (*z*^(*i*)^ → *G*_2_) with probability which depends on its damage level *y*^(*i*)^

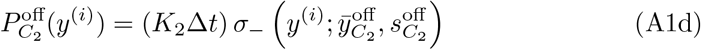

where the positive constants *K*_2_, 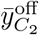 and 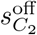 represent, respectively, the maximum rate at which cells leave the G2 checkpoint, the threshold damage level for G2 checkpoint deactivation and the sensitivity of the G2 checkpoint deactivation to damage levels. Eq. (A1d) implies that cells are allowed to re-enter the cell cycle by transitioning to state *G*_2_ only if their damage levels are sufficiently low. By setting 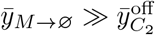, we ensure that cells exiting the G2 checkpoint will successfully undergo mitosis.
- A cell in state *z*^(*i*)^ = *S*, upon completing of DNA synthesis (*x*^(*i*)^ = 2), may arrest due to damage-mediated activation of the G2 checkpoint (*z*^(*i*)^ → *C*_2_) or transition to the next cell cycle phase (*z*^(*i*)^ → *G*_2_) with probabilities

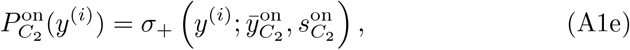

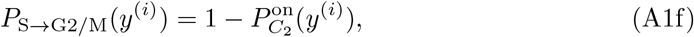

where the positive constants 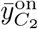 and 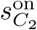 represent, respectively, the threshold damage level for activation of the G2 checkpoint and the sensitivity of G2 checkpoint activation to damage levels. Eq. (A1e) captures the graded damage-mediated activation of the G2 checkpoint (*i.e*., transition to state *C*_2_) that slows a cell’s progression through the G2 phase, to allow damage repair.

#### Algorithm 1

Pseudocode outlining the procedure used to simulate cell proliferation and death. The functions 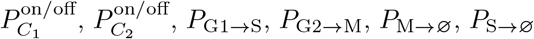 and *P*_G2 →Ø_ are defined by Eq. (A1)-(A2) and M_dNTP_ by Eq. (A3c).

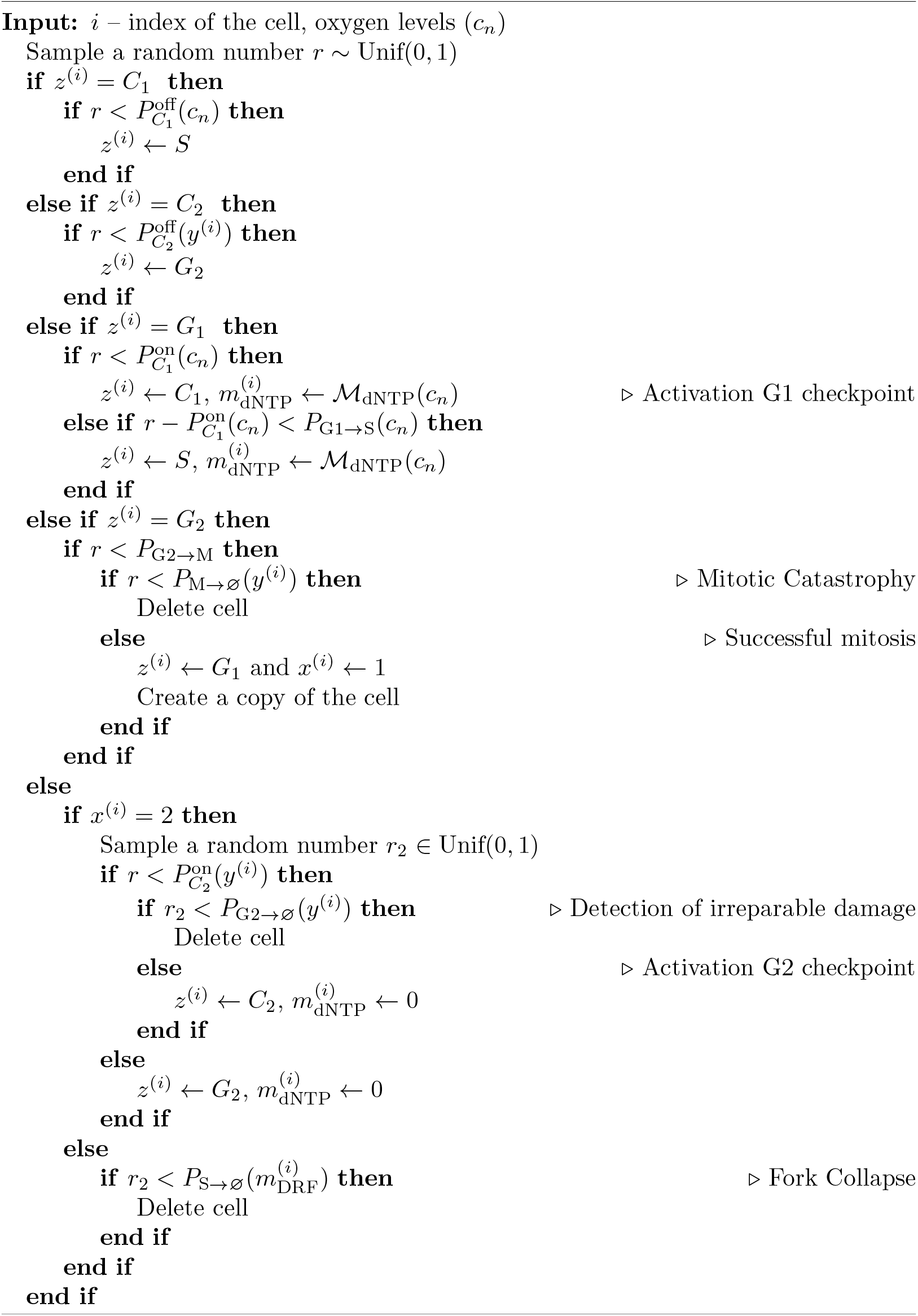

### A.2 Cell fate decision: cell death & division

Based on experimental evidence (see Section 2), we include different forms of replication/cell death, which are cell cycle specific. We assume that cells in the G1 phase (*i.e*., *z* = *G*_1_, *C*_1_) are not sensitive to hypoxia-mediated death, whereas cells in other cell-cycle states, *z*^(*i*)^ ∈ {*S, G*_2_, *C*_2_}, die with probabilities *P*_*z*_(*i*)_*→*∅_ which depend on their damage level *y*^(*i*)^ and/or their DRF expression levels 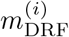 (see Figure 3) in the following way.

- A cell that remains in the S phase may die due to fork collapse with probability

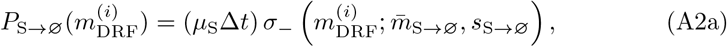

where the positive constants 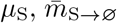 and *s*_S*→*∅_ represent, respectively, the maximum rate of cell death due to fork collapse, the threshold of DRF levels for activation of fork collapse and the sensitivity of fork collapse to DRF levels. In line with the discussion in Section 2, fork collapse is regulated by the intracellular levels of DNA repair factors.
- A cell in state *z*^(*i*)^ = *G*_2_ may attempt mitosis (*i.e*., cell division) with probability

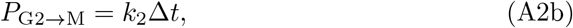

where *k*_2_ *>* 0 is the constant rate at which cells in state *G*_2_ attempt mitosis. During this process, a cell may die via mitotic catastrophe (Matthews et al, 2022) due to the accumulated damage with probability

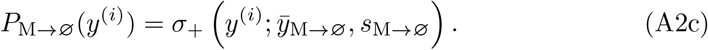 Here, 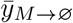 and *s*_*M→*∅_ are positive constants representing respectively the threshold damage level for mitotic catastrophe and the sensitivity of mitotic catastrophe to damage levels. Without loss of generality, the damage levels *y* are rescaled so that 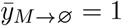.
- A cell that enters the G2 checkpoint, *i.e*., it has transitioned to state *C*_2_, may permanently exit the cell-cycle due to accumulation of irreparable damage with probability

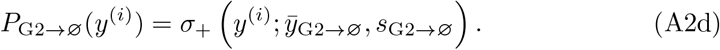 In Eq. (A2d), 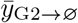 and *s*_G2*→*∅_ are positive constants representing respectively the the threshold damage level for replicative death upon activation of the G2 checkpoint and the sensitivity of replicative death to damage levels. We here take 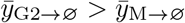 to account for the protective effect of the G2 checkpoint activation. More specifically, for a value of *y*^(*i*)^ a cell’s chances of successfully dividing are increased by activation of the G2 checkpoint. Given the form of Eq. (A2c)-(A2d), the benefit of activation of the G2 checkpoint is maximal for intermediate values of cell damage, and minimal for very low 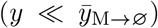 and very high damage 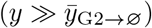 levels.

When a cell dies, it is simply removed from the system. When a cell divides, the original parent cell is removed and two *G*_1_ daughter cells are added. These inherit the values of the state variables *y, m*_dNTP_ and *m*_DRF_ from the parent cell.

### A.3 Modelling the impact of hypoxia on intracellular factors

At each time step, the dynamics of intracellular processes (namely, DNA synthesis, damage repair, dNTP and DRF synthesis/degradation) are simulated within each cell following the procedure detailed in Algorithm 2. Details on the modelling of DNA synthesis and damage repair have been discussed in Section 3.3.1-Section 3.3.2, respectively. We now explain how we account for the impact of hypoxia on dNTP levels (*m*_dNTP_) and DRF levels (*m*_DRF_) in our IB model.

#### A.3.1 Modelling the dynamics of intracellular dNTP levels

Levels of intracellular dNTPs are known to increase upon entry to the S phase (before initiation of DNA synthesis) and to decrease when a cell completes DNA synthesis (Stillman, 2013). In line with these observations, we 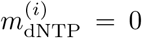 for all cells *I* except those in states *C*_1_ and *S*. Since de-novo production of dNTPs is impaired under low-oxygen, we assume that the change in 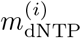 over a time-step Δ*t* satisfies:

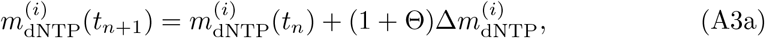

##### Algorithm 2

Pseudocode outlining the procedure used to simulate intracellular processes.

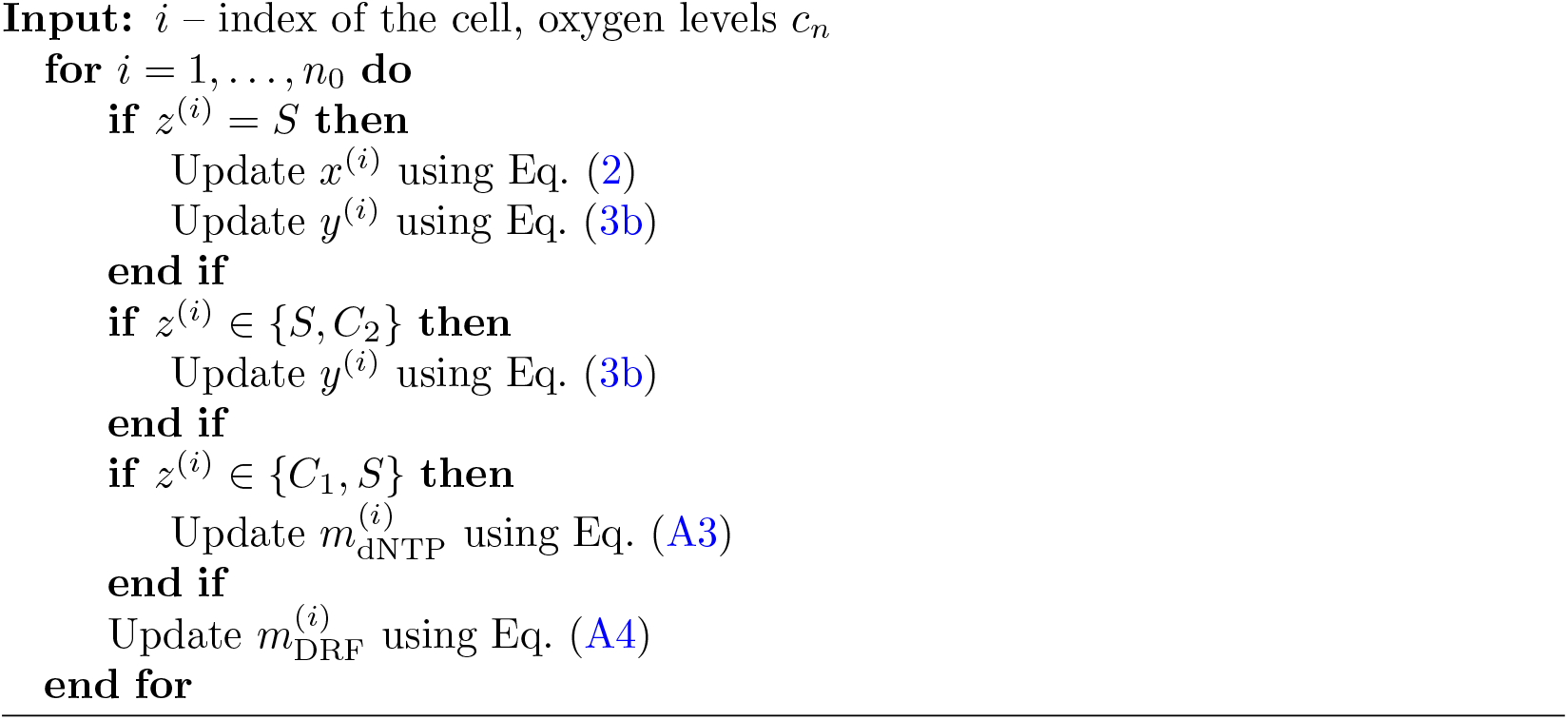

where the multiplicative noise term Θ ∼ 𝒩(0, *σ*) accounts for intercellular variability and we define

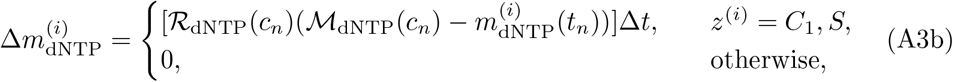

In Eq. (A3b), the function M_dNTP_(*c*) ≤ 1 indicates the baseline expression levels of dNTPs as a function of oxygen conditions, while the positive function ℛ_dNTP_(*c*) indicates the rate at which dNTP levels relax to such baseline values as a function of oxygen. Following (Celora et al, 2022), we account for differences in the dynamics of dNTP levels under physiological and hypoxic oxygen conditions, by setting:

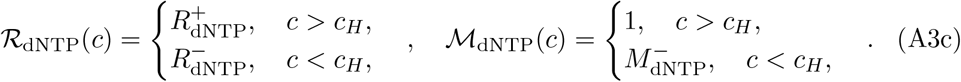

In Eq. (A3c) the positive constants 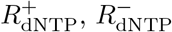 and 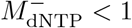 represent respectively the rate of recovery under oxygen-rich conditions, the rate of inhibition in hypoxia and baseline expression levels under hypoxia. When a cell exits the *G*_1_ state, its dNTP levels are set to the baseline value ℳ_dNTP_(*c*).

#### A.3.2 Modelling the dynamics of intracellular DRF levels

We assume that the dynamics of the DNA repair factors do not depend on the cell cycle state (Bindra et al, 2004) and are regulated uniquely by oxygen levels (see Section 2). As for *m*_dNTP_, we model the impact of hypoxia on the expression level 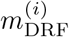 in cell *I* (see Figure 1) by using the following rule to update the DRF expression levels in cell *i* from time *t*_*n*_ to time *t*_*n*+1_:

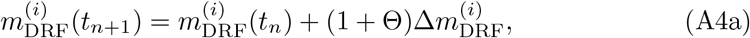

where Θ ∼ 𝒩(0, *σ*) and

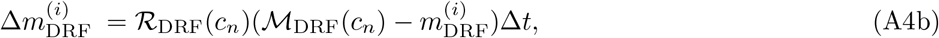

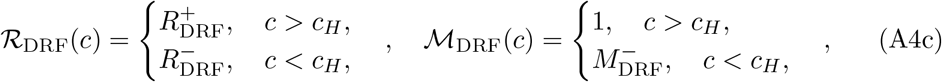

and 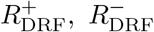 and 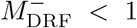 are constant positive parameters representing, respectively, the rate at which DRF levels increase under oxygen-rich conditions, the rate at which DRF levels decrease under hypoxia and the baseline expression levels of DRF under hypoxia. As above, we account for intercellular variability by adding multiplicative noise in Eq. (A4a).

## Appendix B Balanced exponential growth (BEG)

The concept of balanced (or asynchronous) exponential growth (BEG) was first introduced by cell biologists to describe the growth of cell populations (Webb, 1987). This regime describes a situation where the total number of cells, *N*, grows exponentially at a constant rate *λ*^BEG^ (*i.e*., *N* ∝ [exp *λ*^BEG^*t*]) and the distribution of cells in the different phases of the cell-cycle tends to a constant profile, which is independent of how the cells were initially distributed along the cell cycle. Experimentally, BEG is observed in cultures of cells and bacteria at low density (*i.e*., in the absence of competition for space and nutrients).

In an oxygen-rich environment (*i.e*., in physiological conditions), cells have no damage (see Figure 6a), and levels of dNTP and DRF are sufficiently high to guarantee normal cell cycle progress. As a result, cell death is negligible and the number of cells in the checkpoint compartments will tend to zero (*i.e*., no cells in states *C*_1_ and *C*_2_). The only difference in the intracellular state of cells is, therefore, their DNA content, *x*. We investigated BEG for DNA-structured populations in our previous work (Celora et al, 2022) and found that the cell cycle distribution (*i.e*., the fraction of cells in each cell cycle phase: 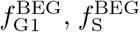 and 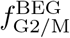) and the growth rate during BEG are related to model parameters as follows

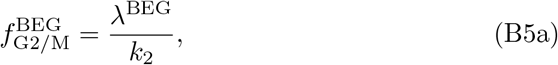

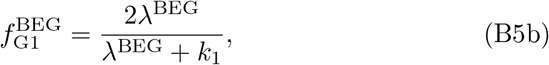

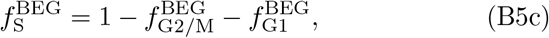

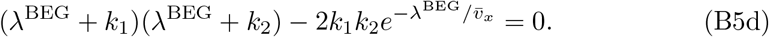

In Eq. (B5), the positive constants *k*_1_, *k*_2_ and 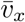 are as in Tables 3-4. The existence and uniqueness of real solutions for Eq. (B5d) are discussed in (Celora et al, 2022). As shown in Figure 7a, the asymptotic behaviour predicted by the IB model agrees with Eq. (B5). Using the results in (Celora et al, 2022), we can also derive the DNA distribution of cells in the S phase

**Table 4:** Summary of the parameters associated with the evolution of intracellular levels of dNTP and DRF used in the simulations that we can estimate from the literature. Typical values are given for the RKO cancer cell line except those taken from other cell lines (indicated with *).

|  | Description | Typical value | Justification (Refs) |
| --- | --- | --- | --- |
| $\Delta t$ | Timestep | 1/50 (hours) | as in Table 3 |
| $\gamma$ | rate at which DNA damage is accumulated in response to replication stress | 0.2 (hours<sup>-1</sup>) | match activation G2 checkpoint in 2/2 cyclic hypoxia (Celora et al, 2022) |
| $\bar{v}_x$ | maximum velocity of DNA synthesis | 0.083 (hours<sup>-1</sup>) | (Celora et al, 2022) |
| $\bar{v}_y$ | rate of DNA repair in physiological levels | 0.3 (hours<sup>-1</sup>) | (in line with values estimated in at low radiation (Liu et al, 2021)*) |
| $R_{\text{DRF}}^-$ | rate of change in the expression of DRFs for $c < c_H$ | 0.13 (hours<sup>-1</sup>) | Figure 14 (Pires et al, 2010a) |
| $R_{\text{DRF}}^+$ | rate of change in the expression of DRFs for $c > c_H$ | 0.05 (hours<sup>-1</sup>) | (≈ 90% recovery in 48 hours (Bindra et al, 2004)*) |
| $R_{\text{dNTP}}^-$ | rate of change in the intracellular levels of dNTPs for $c < c_H$ | 0.3 (hours<sup>-1</sup>) | (Celora et al, 2022) |
| $R_{\text{dNTP}}^+$ | rate of change in the intracellular levels of dNTPs for $c > c_H$ | 0.26 (hours<sup>-1</sup>) | (Celora et al, 2022) |
| $M_{\text{DRF}}^-$ | equilibrium levels of DRFs expression levels in hypoxia | 0.0154 (a.u.) | Figure 14 (Pires et al, 2010a) |
| $M_{\text{dNTP}}^-$ | equilibrium levels of dNTPs levels in hypoxia | 0.06 (a.u.) | minimal rate of DNA synthesis as in (Celora et al, 2022) |

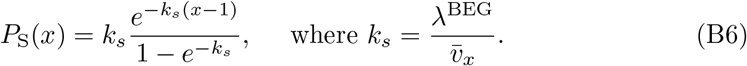

Here *P*_S_(*x*) is the probability that a randomly sampled *S*-type cell has DNA content *x*^(*i*)^ = *x*. In Eq. (B6), the variable *x* has a truncated exponential distribution on the interval [1, 2] with rate *k*_*s*_, *x* ∼ Exp_[1,2]_(*k*_*s*_). In Figure 12, we compare Eq. (B6) with the shape of *P*_S_ estimated from simulations of the IBM model for physiological oxygen levels. The two profiles are in excellent agreement.

**Fig. 12:**
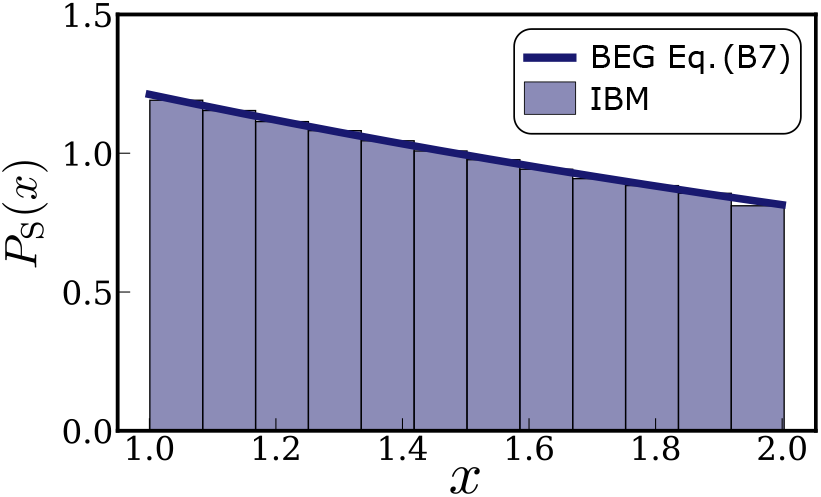
*S*-type cell DNA distribution *P*_S_(*x*) during balance exponential growth. The blue curve represents the theoretical prediction from (Celora et al, 2022) (see Eq. (B6)). The histogram is obtained by direct simulation of the IB model (as in Figure 7). While we simulate the model for 100 hours, we neglect the first 75 hours of the simulation when estimating the DNA distribution; this limits the influence of the initial conditions on the estimated distribution.

We start each simulation with 100 cells. At the beginning of each simulation, an initial state is assigned to each cell as outlined in Algorithm 3. The cell cycle state *z*^(*i*)^ and DNA content *x*^(*i*)^ are assigned by sampling from the BEG distribution. This requires the values of 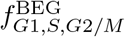 and *k*_*s*_ (see Table 6) and the definition of the cumulative DNA distribution 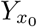.

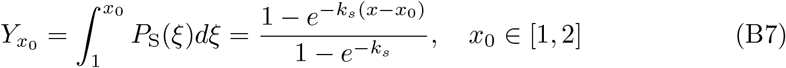

where *P*_S_ is defined by Eq. (B6). The other state variables are initialised to their default physiological values, so that 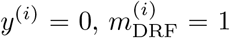, and 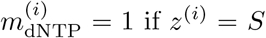 and 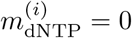 otherwise.

### Algorithm 3

Pseudocode outlining the procedure used to initialise the simulations of our IB model. The function 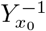 indicates the inverse of the cumulative DNA distribution 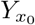 (see Eq. (B7)).

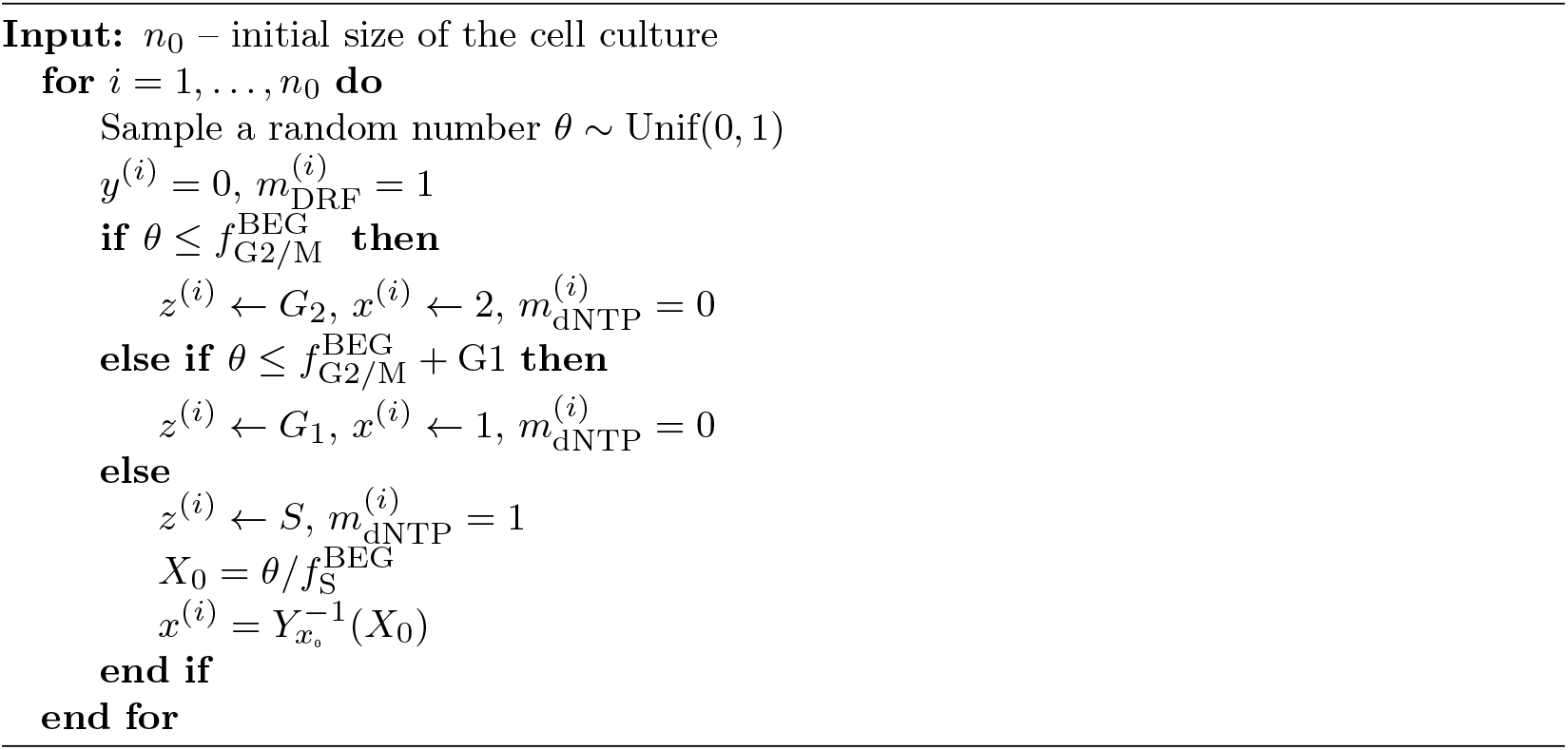

## Appendix C Parameter values

Tables 3-6 contain the model parameters used in the simulations. Where multiple values are indicated, these correspond to cell populations with different damage repair capacities (see Section 3.4.1 and Section 4.3). Where no reference is given, the parameter values have been chosen to produce biologically reasonable behaviour and a justification is given. In particular, the values were chosen to give growth and cell-cycle dynamics in line with experimental & theoretical predictions from (Celora et al, 2022; Bader et al, 2021b).

### C.1 Cell cycle transitions, checkpoint dynamics and cell fate decision

Table 3 lists the value of model parameters associated with cell cycle transition and activation/deactivation of cell cycle checkpoints, *i.e*., Eqs. (A1), and cell-fate decisions, *i.e*., Eqs. (A2).

As mentioned in Section 2, heterogeneity in the regulation of the DNA damage response (DDR) is commonly found in *in vivo* tumours. We model alteration to DDR response by changing model parameters associated with the probability of activation 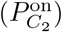 and deactivation 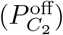 of the G2 checkpoint in response to damage (see Eqs. (A1d)-(A1e)). To model enhanced DDR activation, we decrease 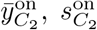 and 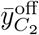 with respect to their default values. To model silencing of DDR signalling, we instead increase 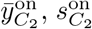 and 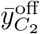 with respect to their default values. Figure 13 shows the profile of 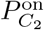 and 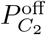 in the three cases: default (DDR^wt^ cells), enhanced DDR activation (DDR^+^ cells) and silenced DDR signalling (DDR^-^ cells).

**Fig. 13:**
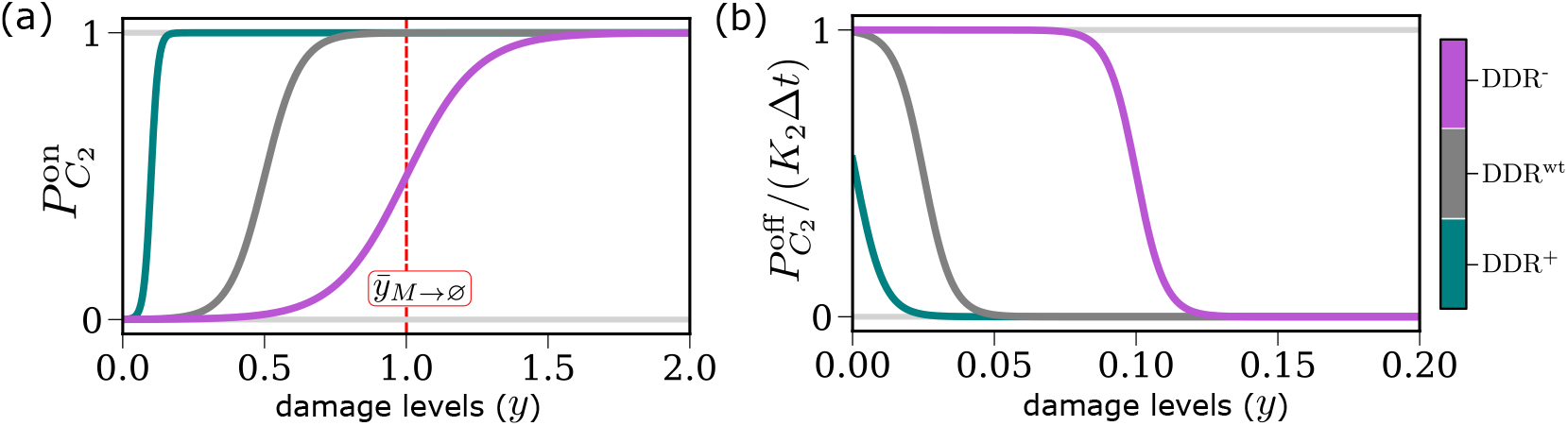
Plots of the profiles of (a) 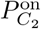 (Eq. (A1e)) and (b) 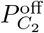 (Eq. (A1d)) for cells with different damage repair capacity. The red dotted line in Figure 13a indicates the curve 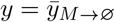, where 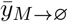 corresponds to the threshold damage level for mitotic catastrophy. Parameter values are as indicated in Table 3.

### C.2 Intracellular dynamics

Table 4 lists the value of model associated with intracellular dynamics, *i.e*., Eqs. (2)-(3) and (A3)-(A4). We estimate the parameters associated with the expected evolution of repair protein expression levels, *m*_DRF_(*t*) (see Eq. (A4)), using the data from (Pires et al, 2010a) on the time-evolution of expression levels of the DNA repair protein RAD51 in RKO cells in constant hypoxia. While in our model *m*_DRF_(*t*) corresponds to the expression of multiple different damage repair proteins, we assume they behave similarly in response to hypoxia and use RAD51 dynamics as a prototypical response. As shown in Figure 14, the expression levels relative to physiological conditions show a clear trend: the expression levels decrease monotonically as the period of exposure to hypoxia increases. The profile can be fitted to an exponential function (see the continuous curve in Figure 14) justifying the functional form chosen for *m*_DRF_(*t*) (see Eq. (A4b)). In this case, model parameters can be inferred directly by interpolation of the data points with an exponential function 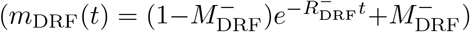. We could not find similarly detailed data for RKO cells to estimate 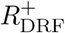, *i.e*., the rate at which protein expression levels are restored upon reoxygenation. Estimates of 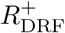 were informed by Western-blot data from Bindra et al (2004), which were collected from experiments on other cell lines.

**Fig. 14:**
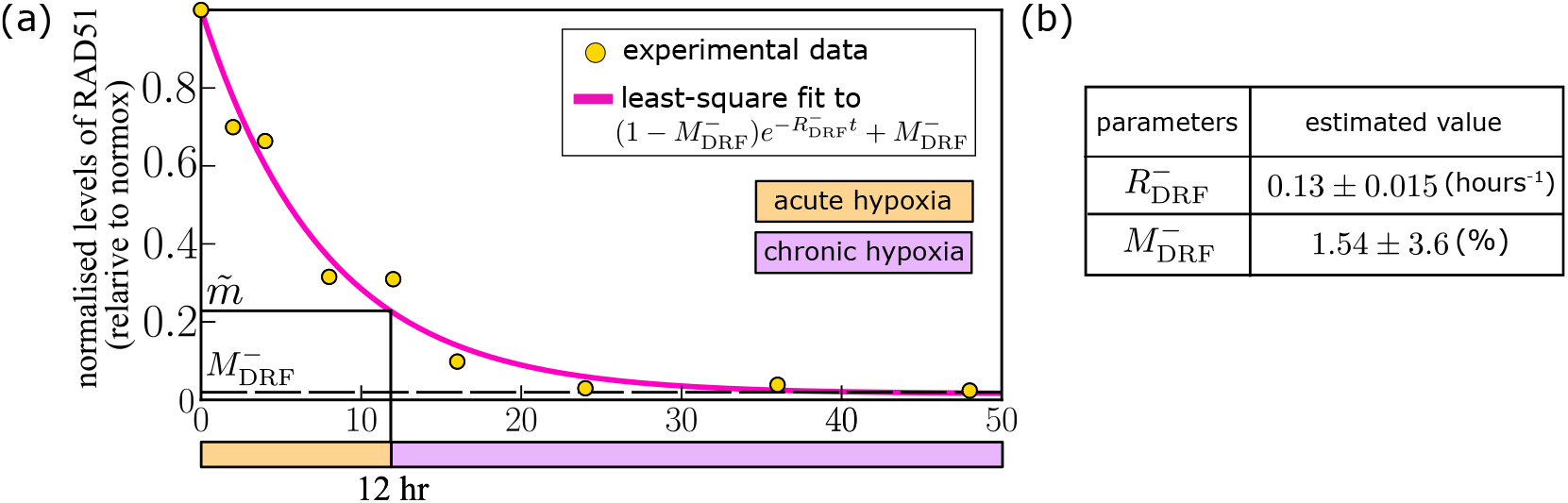
(a) Data from (Pires et al, 2010a) on the time-evolution of expression levels of the DNA repair protein RAD51 in constant hypoxia (yellow dots). The pink curve indicates the value of the function 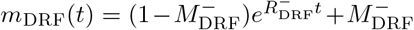 (obtained by solving the deterministic version of Eq. (A4) starting with *m*_DRF_(0) = 1 and setting *c*(*t*) ≡ *c*_*−*_ *< c*_*H*_ for all *t*) for parameters values as specified in the panel (b). (b) Estimates of the parameters 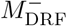 and 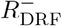 are obtained by fitting the function 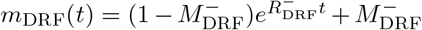 to the experimental data using the `curve_fit` function in the *SciPy* library in Python. For each parameter, we indicate the computed 67% confidence interval.

Using the data in Fig. 14, we find that *m*_DRF_ drops to 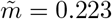 after 12 hours in hypoxia. As discussed in Section 2, experimentally it is observed that, when exposed to hypoxia for more than 12 hours, cells become sensitive to the collapse of replication forks and repress repair mechanisms. Accordingly, we require that the death rate for cells in the S phase is half of its maximum value when cells are exposed to constant hypoxia for 12 hours, *i.e*., we set 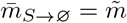 in Eq. (A2a).

### C.3 Oxygen dynamics in the chamber

Table 5 lists the value of model parameters used to simulate the evolution of oxygen levels within the oxygen chamber for different experimental setups (see Eqs. (4)).

**Table 5:**
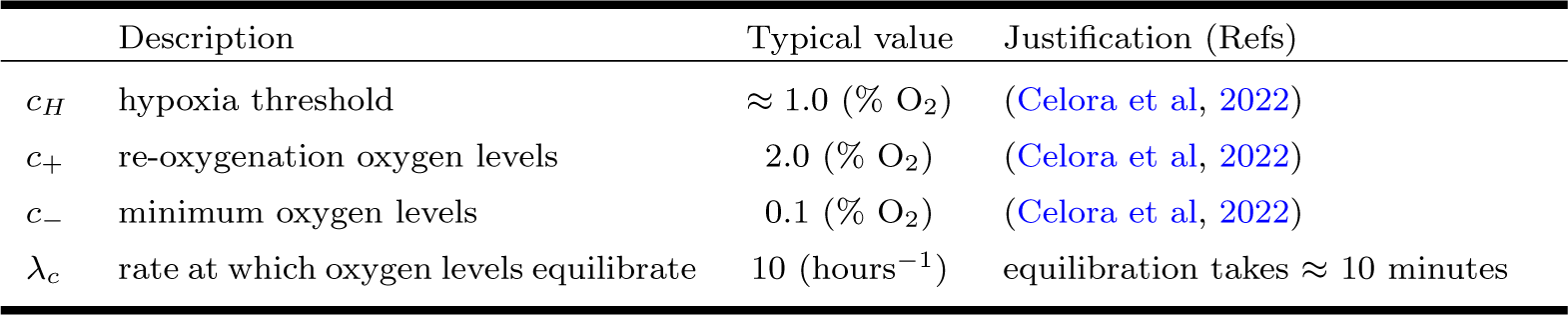
Summary of the parameters associated with the evolution of oxygen levels within the chamber used and the values used in the simulations (see Eqs. (4)).

**Table 6:** Summary of the parameters associated with the initialisation of the numerical simulations (see Section B).

|  | Description | Typical value | Justification (Refs) |
| --- | --- | --- | --- |
| $f_{G1}^+$ | initial fraction of cells in the G1 phase | 29 (%) | (Celora et al, 2022) |
| $f_S^+$ | initial fraction of cells in the S phase | 56 (%) | (Celora et al, 2022) |
| $f_{G2}^+$ | initial fraction of cells in the G2 phase | 15 (%) | (Celora et al, 2022) |
| $k_s$ | rate at which cell leave the S phase in the BEG regime | $\approx 0.4$ | see Eq. (B6) (Celora et al, 2022) |

### C.4 Initial conditions: BEG

Table 6 lists the value of model parameters used to initialise the numerical simulations (details can be found in Section B).

## Appendix D Initial cell cycle state influences cell survival

Figure 15 shows the estimates of survival obtained via numerical simulations of clonogenic assays (see Section 3.4.2) stratified by the cell cycle phase of the initially seeded cells. We compare results for four cyclic hypoxia conditions; namely, (4,5)-, (7,5)- and (11,5)-cyclic hypoxia. For oxygen conditions where overall population survival is high (*e.g*., (4,5)-cyclic hypoxia) or low (*e.g*., (11,5)-cyclic hypoxia), there is no significant difference in cell survival for cells starting in different cell cycle phases. Larger deviations are observed for conditions where the overall population survival 𝒱 ≈ 0.5 (*e.g*., (7,5)-cyclic hypoxia). In this example, progenitor cells initially in the late stages of the cell cycle –*i.e*. in the S and G2/M phases – are more and more likely to survive. As the duration of 𝒯_*R*_ decreases and we approach conditions analogous to constant hypoxia (*e.g*., (11,0.5)-cyclic hypoxia), we observe an even stronger correlation between cell survival and the initial cell cycle phase of the progenitor cell. Overall, the results presented in Figure 15 highlight the importance of deconvoluting cell cycle-specific sensitivities when accessing survival in toxic environments.

**Fig. 15:**
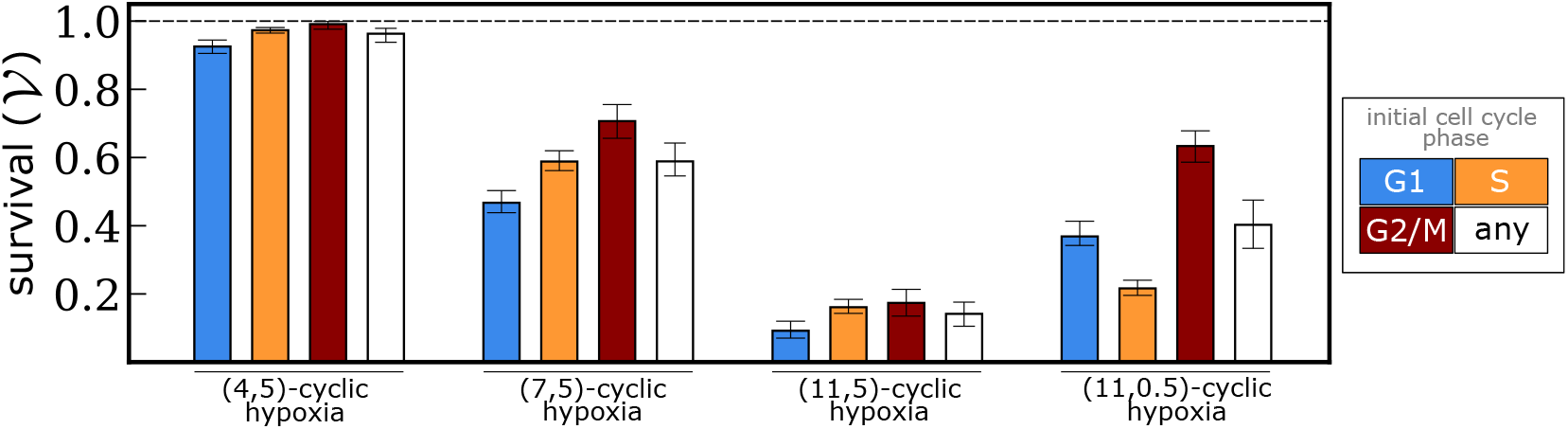
Characterising the impact of the initial cell cycle state of progenitor cells on survival. Barplots indicate the probability of survival, 𝒱 estimated via simulation of clonogenic assay experiments (see Section 3.4.2) stratified by the cell cycle phase of the progenitor cell. We compare the results for four cyclic hypoxia conditions: (4,5), (7,5), (11,5), and (11,0.5)-cyclic hypoxia. Values of the parameters are as in Figure 7.

## Appendix E Long-term damage distribution in serial passage experiments

In Section 4.3, we claim that, during serial passage experiments in cyclic hypoxia, the distribution of damage within the population does not converge to a stationary distribution at long times; rather it fluctuates with the oxygen levels, and it eventually settles to a time-periodic function whose period coincides with the interval between passages of the cell population. We here present additional results in support of our claim.

Figure 16 illustrates the time-evolution of the median and IQR of the damage distribution in the population during serial passage experiments (see Section 3.4.3) for cells exposed to (4,5)-cyclic hypoxia. We report the simulated time-evolution for both the co-culturing (see Figure 16a) and control experiments (see Figure 16b). The results show that the damage distribution statistics do not settle to stationary values but rather they fluctuate with the oxygen levels. At long times, the distribution converges to a periodic function, whose period is equivalent to the interval between contiguous passages of the cell population (here 45 hours).

**Fig. 16:**
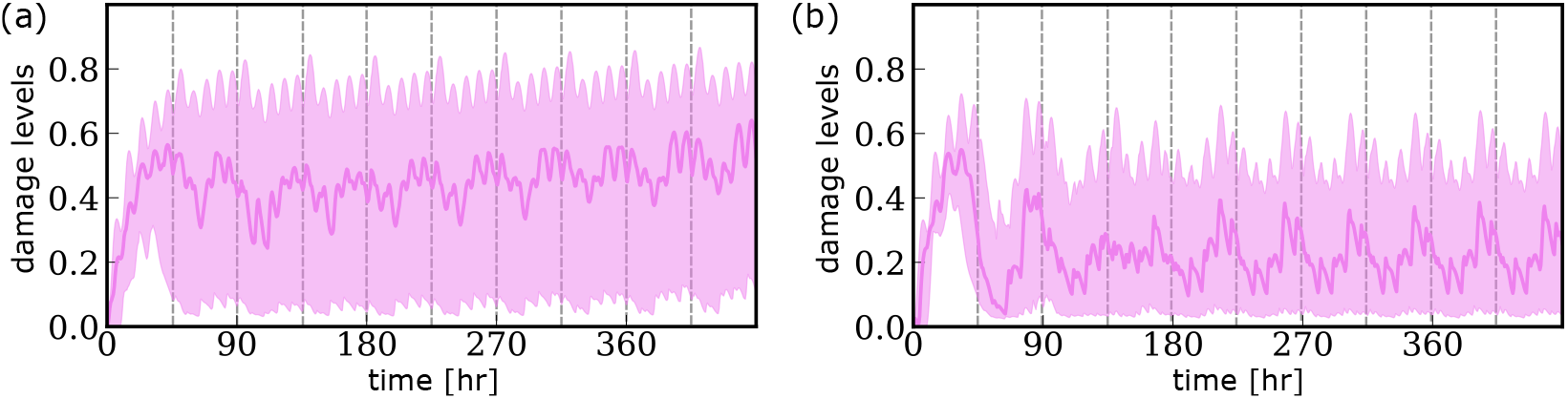
Plots of the time-evolution of the damage distribution during simulations of serial passage experiments in (4,5)-cyclic hypoxia for (a) co-culture and (b) control conditions. We indicate the median (see violet curve) and the interquartile range (see shaded area) of the damage distribution extracted from the simulations from Figure 11. The vertical dotted lines indicate the time at which cells are passed (*i.e*., replated).

